# Novel strategy for treating neurotropic viral infections using hypolipidemic drug Atorvastatin

**DOI:** 10.1101/639096

**Authors:** Suvadip Mallick, Surajit Chakraborty, Bibhabasu Hazra, Sujata Dev, Sriparna Mukherjee, Masood Ahmad Wani, Anirban Basu

**Affiliations:** National Brain Research Centre, Manesar, Haryana, 122052, India

## Abstract

Chandipura virus (CHPV) and Japanese Encephalitis Virus (JEV) are known to infect neurons followed by their successful propagation. Increased incidences of central nervous system invasion by the abovementioned viruses have been reported in case of children and elderly thus culminating into severe neurological damage. Literature suggests induction of endoplasmic reticulum (ER)-stress related proteins upon CHPV and JEV infection which help promote viral reproduction. Since earlier studies underscore the pleotropic role of atorvastatin (AT) in neuroprotection against flaviviruses like Hepatitis C and dengue, it was hypothesized that AT might also act as a neuroprotective agent against RNA viruses like CHPV and JEV. AT-mediated antiviral activity was evaluated by assessing survivability of virus-infected mouse pups treated with the drug. Balb C mice were used for *in vivo* experiments. Neuro2A cell line was used as the model for *in vitro* experiments. Cells subjected to AT treatment were infected by CHPV and JEV followed by evaluation of ER stress-related and apoptosis-related proteins by immunoblotting technique and immunofluorescence microscopy. Interaction of host protein with viral genome was assessed by RNA-Co-IP. AT treatment exhibited significant anti-viral activity against CHPV and JEV infections via hnRNPC-dependent manner. Viral genome-hnRNPC interaction was found to be abrogated upon AT action. AT was also observed to reduce secretion of proinflammatory cytokines by the neurons in response to viral infection. Moreover, AT treatment was also demonstrated to reduce neuronal death by abrogating virus-induced miR-21 upregulation in hnRNPC-dependent fashion. This study thus suggests probable candidature of AT as antiviral against CHPV and JEV infections.

## Introduction

Incidences of Acute Encephalitis Syndrome (AES) resulting from infection by neurotropic RNA viruses like Japanese Encephalitis Virus (JEV) and Chandipura Virus (CHPV) have been reported in India on an annual basis (1). Both JEV and CHPV are arboviruses differing in terms of genetic material. JEV possesses a single positive-stranded RNA as genetic material, while CHPV is a negative-stranded RNA virus (2, 3). JEV is being reported to mainly target central nervous system (CNS) of children. Since CNS of infants undergo a lot of physiological changes as development progresses, any insult to neuronal physiology may lead to severe perturbation of neurocognitive abilities thus leaving the survivors of JEV-induced encephalitis with motor deficits, cognitive and language impairments and learning difficulties (4, 5). Whereas, although infection due to CHPV leads to acute encephalopathy, no report till date documents incidences of neurological sequel experienced by survivors of CHPV-induced encephalitis (5). Evidences suggest neuronal death following JEV or CHPV infections by two distinct mechanisms. In addition to direct viral propagation-mediated neuronal result, microglial activation-triggered inflammatory storm also has been reported to culminate into bystander death of neurons (6, 7). No effective antivirals exist against these virus infections till date, and thus therapy administered to patients suffering from aforementioned viral infections are aimed at reducing severity of signs and symptoms. This therefore warrants the need for extensive investigation for discovery of compounds helping in combating JEV and CHPV infections. Owing to the great amount of time and painstaking labor behind new drug discovery, repurposing of drugs has revolutionized drug discovery as an alternative to new drug discovery pipeline associated with high cost and high failure rates. Thorough research is being carried out all over the world in search of more candidate drugs suitable for repurposing against diseases for which any specific treatment is not yet present. For instance, a recent study by Kumar *et al*., 2015 provides evidence for role of minocycline in ameliorating Acute Encephalitis syndrome (AES)-associated mortality rate, thus warranting large cohort studies in order to evaluate minocycline’s efficacy as a potent drug against acute encephalitis syndrome (8).

Statins are chiefly advertised owing to their ability to reduce serum cholesterol levels (9, 10). Statins have also been reported to be successfully used as preventive therapeutic strategy against neurodegenerative diseases: e.g., in the settings of cerebrovascular accidents (11), Parkinson’s disease (12), Alzheimer’s disease (13), and multiple sclerosis (14). A recent report also demonstrates effectiveness of statins in treating traumatic brain injury (15). There have been several reviews which thoroughly document clinical and experimental data thus discussing both pros and cons regarding use of statins against neurodegenerative diseases (16, 17, 18, 19, and 20). Atorvastatin belonging to the class of hydroxymethylglutaryl coenzyme-A reductase (HMG-CoA-reductase) inhibitors is a well-tolerated drug aimed at lowering serum lipid levels. Stability of atorvastatin has been observed to increase upon undergoing lactonization, thus resulting in the ability of the same in crossing blood-brain barrier (BBB). *In vivo* lactonization of atorvastatin has been demonstrated by Jacobsen *et al*, upon administration of atorvastatin as acid (21). Moreover, recent reports support the use of atorvastatin for HIV-infected children with hyperlipidemia (22).

At the first place, study by Fedson *et al* pointed towards the use of statin in amelioration of pandemic mortality (23). Since the first report, use of stains as preventive medicine aimed at treating infection-induced cellular damage has been documented already by several studies (23, 24). A considerable body of research demonstrates pleotropism exhibited by statins comprising of modulation of cellular signaling pathways, inflammatory gene transcription and anti-inflammatory and immunomodulatory characteristics. Statins have also been shown to abrogate sepsis and infection-associated inflammation independent of their lipid-lowering potential (25). Study by Sun *et al* also underscores the importance of statins in negatively interfering with key signaling pathways in response to viral infection (26). Effects of statins in limiting inflammation and cellular toxicity associated with influenza infection thus suggest use of statins as an alternative in treating the same (27). Study by Haidari *et al* reports inhibitory effect of statin administration upon influenza virus propagation in Madin-Darby canine kidney (MDCK) cells. The abovementioned action of statins was observed to act via downregulation of Rho/Rho kinase pathway activity. Finally, the same study also demonstrates that atorvastatin also reduced C57BL/6 mice mortality and viral titer in the lung upon infection with H3N2 and H1N1 (28). Investigations also point towards the efficacy of atorvastatin treatment in reducing abundance of proinflammatory cytokines like TNF- and IL-6 in response to H1N1 infection of Crandell feline kidney (CrFK) cells (29). These abovementioned studies thus warrant further future research aimed at investigating the full potential of statins in acting as a supplementary therapy alongside with conventional therapeutic regimes in controlling the cytokine overproduction in virus infections. In addition to its immunomodulatory action, atorvastatin treatment was shown to be as effective as oseltamivir and amantadine in decreasing viral titer and increasing cell viability upon H1N1 infection in MDCK cells (30). Atorvastatin has also been found to limit early brain injury by exhibiting anti-apoptotic effects and down-regulation of ER stress in a rat model of subarachnoid hemorrhage (31), and also suppresses the development of abdominal aortic aneurysm in mouse model (32).

CHPV and JEV are single-stranded RNA viruses, and are being reported to infect neurons thus culminating into neuronal death. Provided with the wide array of results demonstrating importance of atorvastatin in combating virus infections via modulation of inflammation, metabolism, we hypothesize that atorvastatin might play role in altering CHPV and JEV propagation in neurons thus effecting survivability in a positive manner. In our present study, we describe that atorvastatin treatment confers effective neuroprotection against CHPV and JEV infection *in vivo* and *in vitro* by lowering CHPV and JEV multiplication. Furthermore, atorvastatin treatment also curtailed inflammatory response upon invasion of viruses into the brain, thus enhancing survivability of the infected mouse pups. Evidences have also been provided in the support of the fact that atorvastatin treatment down regulated ER stress-induced heterogeneous nuclear ribonucleoprotein C1/C2 (hnRNP C1/C2) proteins upon infections. To investigate whether regulation of hnRNPC1/2 acts as the pivotal mechanism by which atorvastatin exerts its action upon viral infection and neuronal apoptosis, hnRNPC abundance was subjected alterations *in vitro* and the effect of atorvastatin treatment was examined in the context of virus infections. Our study also demonstrates the role of atorvastatin in negatively regulating interaction of hnRNPC and viral genomes *in vitro* and *in vivo*, thus providing a possible mechanism of the drug action upon CHPV and JEV infections. Since miRNAs are reported to play crucial roles in shaping numerous physiological and pathological responses, role of atorvastatin in reprogramming expression of apoptosis-associated miRNA was also investigated. We have shown that atorvastatin treatment reduced infection-induced miR-21 upregulation, which is normally known to promote cellular apoptosis. Suppression of miR-21 activity has been demonstrated to reduce cellular death in a PDCD4-dependent manner.

## Materials and Methods

### Ethics statement

All animal experiments performed were approved by the animal ethics committee of National Brain Research Centre (Approval no.: NBRC/IEAC/2017/128, NBRC/IEAC/2018/146). Animals were handled as per strict guidelines defined by the Committee for the Purpose of Control and Supervision of Experiments on Animals (CPCSEA), Ministry of Environment and Forestry, Government of India. Human autopsy tissue samples were collected from the Human Brain Bank, NIMHANS, Bangalore as per institutional ethics and confidentiality of the subjects.

### Virus and cells

CHPV (strain no. 1653514), isolated from a human patient at Nagpur, India, 2003, was a kind gift from Prof. Dhrubajyoti Chattopadhyay (Amity University, New Town, Kolkata). Suckling BALB/c mice were used for the propagation of GP78 strain of JEV. Following intracranial administration of JEV, the mice pups were carefully monitored for signs of infection. Following the appearance of signs, brains of infected pups were harvested and subjected to titer measurement as mentioned earlier (33). Titer of CHPV and JEV were found to be 10^9^ PFU/ml and 10^8^ PFU/ml respectively. Neuro2A cells, Porcine stable kidney (PS) cells (both obtained from National entre for Cell Science, India) and Vero E6 cells (kind gift from Prof. Debi P. Sarkar, Delhi University, South campus,) were grown at 37°C in Dulbecco’s modified Eagle medium supplemented with 3.5% sodium bicarbonate, 10% fetal bovine serum and penicillin-streptomycin.

### Animal treatment

We used 10-days old BALB/c mouse pups irrespective of sex for the experiments. While the experiments were being performed, mice pups were housed with their mother for the purpose of feeding. The animals were randomly divided into four groups: Control mock-infected group (MI), only atorvastatin-treated group (MI+AT), Infected groups (CHPV or JEV) and atorvastatin-treated group (CHPV+AT or JEV+AT).The CHPV-infected group was administered with approximately 1.25×10^5^ viral particles suspended in 50 µl of phosphate buffer saline (PBS) through intraperitoneal route.. Mice pups belonging to the JEV-infected group were similarly infected with JEV (approximately 1.25×10^4^ particles) suspended in 50 µl of PBS. Atorvastatin Calcium trihydrate (kind gift from Sun pharmaceuticals, erstwhile Ranbaxy laboratories) was dissolved in DMSO was further diluted in 1X sterile PBS and administered intraperitoneally at a dosage of 5mg/kg-body weight once followed by 3 hours of infection. AT treatment was continued for 4 and 7 days in CHPV and JEV-infected group respectively. CHPV and JEV-infected groups succumbed to infection by 3-4 days and 6-7 days post-infection respectively followed by their sacrifice and collection of infected brain samples. The harvested infected-brain samples were stored at −80°C for further processing. For immunohistochemical experiments, brains were collected following transcardial perfusion with ice cold 1X PBS followed by tissue fixation using 4% paraformaldehyde (PFA). To decide whether atorvastatin has its role in delaying the manifestation of clinical symptoms in the treated groups or not, random animals were left to check the survival in the treated groups.

### Infection and treatment to Neuro2A cells

Neuro2A cells were cultured till 60% to 70% confluence was achieved, followed by differentiation in serum free medium. Cells were divided into six groups: Control mock-infected group (Control), only atorvastatin-treated group, where mock-infected cells were treated with two different doses of AT (1 µM and 2 µM), virus-infected group (CHPV or JEV) and virus-infected cells treated with AT (CHPV+1 µM AT and CHPV+2 µM AT or JEV+1 µM AT and JEV+2 µM AT). Cells were infected with CHPV and JEV at a multiplicity of infection (MOI) of 0.1 and 1 respectively. Following 2 hours of infection, cells were washed thoroughly with sterile 1x PBS to remove the unwanted attached virus particles and atorvastatin was added to the maintenance medium. Cells were then incubated at 37°C for different time points as per experimental paradigms. Neuro2A cells pretreated with Thapsigargin (1 µM) for 2 hours were then maintained in AT-containing media followed by their collection after 24 hours for further studies.

### Primary neuron culture

Mouse cortical neurons were cultured as per previously-mentioned protocol (34). Briefly, 2-day old BALB/c mouse pups’ brain cortex was isolated in calcium and magnesium-free tyrode solution, under a dissecting microscope and sterile conditions. Trypsin and DNase enzymes were used to homogenize cortex to make single cell suspension which was made to undergo filtration through a 127 μm pore size nylon mesh (Sefar) to eliminate cellular debris. The filtrate was then subjected to centrifugation and the supernatant was discarded. The pellet collected was resuspended in serum containing media. Cells counted using hemocytometer were seeded in equal numbers onto poly-D-lysine (Sigma, USA)-coated plates and maintained in neurobasal medium containing 2 mM L-glutamine, 1% glucose, 5% FBS, horse serum and penicillin-streptomycin. After two days, aforementioned serum-containing complete media was removed to inhibit glial growth followed by addition of N2 and B27 supplement-containing neurobasal media. 20 μM Ara-c (cytarabine) was added to the culture one day prior to viral infection to eliminate the astrocytes. Primary neuronal culture was infected with CHPV and JEV and then treated with atorvastatin (2 µM) similarly to aforementioned procedure for infecting and treating Neuro2A cells.

### RT-PCR and qRT-PCR

Following isolation of total RNA from N2A cells and mouse brain samples using Tri reagent (Sigma-Aldrich), 250 ng of RNA was reverse transcribed with the help of Verso cDNA synthesis kit (Thermo Fisher Scientific). The sequences of the PCR primers used in this study are as follows: CHPV forward primer 5ʹ-GATCGCGGAGTGGTAGAATATC-3ʹ, CHPV reverse primer 5′-GAAATCAGCCATGTGTTGTCC-3′; JEV forward primer 5ʹ-CAGGGAAGAGATCAGCCATTAG-3ʹ, JEV reverse primer 5ʹ-GGAGCATGTACCCATAGTGAAG-3ʹ; GAPDH forward primer 5ʹ-ATGGCAAGTTCAAAGGCACAGTCA-3ʹ, GAPDH reverse primer5ʹ GGGGGCATCAGCAGAAGG-3ʹ.The thermocycling conditions used for polymerase chain reactions were 95°C for 30 seconds, 54°C for 45 seconds, 68°C for 1 minute. Optimum number of cycles comprising of aforementioned conditions were 35. The amplified products were then resolved using 1% agarose gel. Ethidium Bromide was used in staining gel and photographed. For quantitative determination of PDCD4 mRNA and miR-21 expression, qRT-PCR analysis was performed. qRT-PCR analysis of PDCD4 was performed with Power SYBR Green PCR Master Mix (Applied Biosystems) with gene-specific forward and reverse primers. 5´-ATGGATATAGAAAATGAGCAGAC-3´ and 5´-CCAGATCTGGACCGCCTATC-3´ sequences were used as forward and reverse primers for the quantification of PDCD4 mRNA. Thermocycling conditions used for determination of PDCD4 abundance has been mentioned earlier (35). Isolation of miRNA and cDNA preparation were performed as described earlier (35). Sequences 5ʹ-UAGCUUAUCAGACUGAUGUUGA-3ʹ and 5ʹ-UAGCUUAUCAGACUGAUGUUGA-3ʹ were used as forward primers in qRT-PCR analysis of human and mouse miR-21 respectively. Human autopsy tissue samples were collected from the Human Brain Bank, NIMHANS, Bangalore, as per institutional ethics and confidentiality of the subjects. miRNA isolation from human brain sections was performed using methods as described previously (35). Expression data for snRNA SNORD68 was used as a normalization control. The thermal cycler QuantStudio 5 (Applied Biosystems) was used for qRT-PCR, and the data were analyzed with the QuantStudio design and analysis software.

### miRNA inhibitor transfection

miR-21 inhibitor and its negative control were purchased from Qiagen. Cells were seeded in six-well plates and transfected with miR-21-inhibitor or inhibitor-control using Lipofectamine 2000 (Invitrogen, Carlsbad, CA, USA) following manufacturer’s instructions. Following 24 hours of transfection, cells were infected with CHPV or JEV for specific time periods. Expression of miR-21 and its target gene was then studied using qRT-PCR and western blotting procedure.

### Cytokine bead array

BD cytometric bead array (CBA) mouse inflammation kit (Cat. No. 552364) was used to measure abundance of inflammatory cytokines in mouse brain lysates and the assay was performed according to the manufacturer’s instructions. BD FACS Calibur (Becton Dickinson, San Diego, CA, USA) was used for analyzing processed samples.

### Plaque assay

Virus titers from *in vitro* and *in vivo* samples were evaluated by plaque assay as described in (36). In short, CHPV-infected and atorvastatin-treated infected-brain samples were homogenized followed by virus isolation. Next, serial dilutions of purified virus-containing solutions were used to infect monolayer of VeroE6 cells. Following infection for 2 hours, VeroE6 cells were washed with sterile 1X PBS and were covered with solution containing 1% low melting-agarose, 1X MEM and FBS, and was solidified. Infected-VeroE6 monolayer covered with agarose overlay was incubated for 3 days at 37°C in the presence of 5% CO_2_. Overlay was then removed and cells were stained with crystal violet for demonstration of plaques. Supernatant from CHPV-infected and AT-treated virus-infected Neuro2A cells was collected and subjected to plaque assay using abovementioned protocol. JEV titer of *in vivo* as well as *in vitro* samples was evaluated using plaque assay. For measurement of JEV titer, PS cell lines were used which were incubated for 96 hours following solidification of agarose overlay.

### Cytotoxicity and Reactive oxygen species assay

In order to determine optimum dosage of AT, AT-mediated cytotoxicity upon Neuro2A cells was measured by 7-AAD-staining. Following staining as per manufacturer’s protocol (BD Pharmingen), cells were analyzed using FACSVerse^TM^ (BD Biosciences) flow cytometer. Since AT was reconstituted in DMSO, toxic effects of DMSO upon cell viability and virus propagation was estimated by treating uninfected and virus-infected Neuro2A cells with DMSO. In brief, DMSO-treated samples were subjected to immunoblotting followed by analysis of abundance of viral proteins and cleaved fragment of pro-caspase-3. The generation of ROS in Neuro2A cells was measured by DCFDA-cellular ROS assay kit (DCFDA; Sigma). Uninfected and infected cells were treated with DCFDA followed by incubation in the dark for 30 minutes. Cells were then subjected to centrifugation and the pellet was collected. Pelleted cells were resuspended in chilled 1X PBS. The intracellular fluorescence was then measured by flow cytometry using FACS Calibur (BD Biosciences, San Diego, CA, USA). Cells pre-treated with N-acetyl L-cysteine (scavenger of ROS; Sigma) was included in the experiment as negative-control.

### Immunoblotting

Proteins isolated fromNeuro2A cells and individual mouse brain lysates across all groups were resolved by SDS-PAGE. Concentrations of protein samples obtained from both *in vivo* and *in vitro* experiments were estimated by bicinchoninic (BCA) protein assay. After separation of proteins by SDS-PAGE, proteins were transferred onto nitrocellulose membrane. Following transfer, membrane was treated with 5% non-fat milk solution to avoid non-specific binding of primary antibodies. The membrane was then incubated in the presence of primary antibodies against CHPV glycoprotein-G, nucleoprotein-N, matrix protein-M (Bharat Biotech; 1:1000), JEV non-structural protein 3 (Genetex; 1:10000), mouse cleaved-Caspase 3 (Cell Signaling, 1:2000), heterogeneous nuclear ribonucleoprotein C1+C2 (Abcam, 1:2000), GRP78 Bip (Abcam, 1:2000), prohibitin (Abcam, 1:2000), Flag-tag (Sigma-Aldrich, 1;1000) and β-actin (Sigma-Aldrich; 1:10,000) overnight at 4 °C with gentle shaking. β-actin was used as loading-control. After vigorous washing with 1X TBST solution (containing tween20), membrane was incubated with respective HRP-conjugated secondary antibodies (Vector Laboratories, USA; 1:5000). Blots were then processed for development chemiluminescence reagent ECL (Millipore, CA, USA) followed by image capture using UVITECH imaging system (Cambridge) (Millipore, CA, USA).

### Immunofluorescence

Brain sections were subjected to permeabilization using 0.1% Triton X-100-containing 1X PBS solution. Following the above mentioned step, sections were incubated in a blocking solution at room temperature for 1 hour. Sections were then incubated overnight along with anti-CHPV (1:200, Bharat Biotech) or anti-JEV NS3 (1:500, Genetex) and anti-hnRNP C1+C2 (1:500, Abcam) antibodies at 4°C. Post-primary antibody incubation, washing was performed, followed by incubation with respective Alexa-Fluor 488 or Alexa-Fluor 594– conjugated secondary antibodies (1:1500, Molecular Probes) for 1 hour. Finally, the sections were mounted with 4′, 6-diamidino-2-phenylindole (Vector Laboratories Inc.) and observed under Zeiss Apotome microscope at the specified magnification. CHPV or JEV-infected primary cortical neuronal cells treated without or with AT atorvastatin for 12 hours and 24 hours respectively, were fixed using 4% PFA, and then processed similarly for immunofluorescence analysis.

### RNA Co-IP and sequencing

Interaction of CHPV and JEV RNA with hnRNPC was analyzed by RNA Co-IP method as described earlier (37). Briefly, Uninfected and infected Neuro2A cells with or without AT treatment were treated with 1% formaldehyde for 10 minutes at room temperature (RT). 0.25 M glycine (pH 7.0) was added to this solution for inducing crosslinks between RNA and proteins. Cells were then subjected to sonication thus releasing the cross-linked RNA-Protein complex. The cell lysate was incubated with superparamagnetic beads covalently linked to recombinant protein G (Dynabeads protein G Novex, Life technologies, USA) to obtain the precleared lysate. Beads were coated with anti-mouse hnRNPC (Abcam) and allowed to incubate with the precleared lysate for 30 minutes at RT under shaking condition.

These beads were then washed with antibody binding and washing buffer followed by elution of the complex using the elution buffer supplied with the kit. RNA was extracted from eluate using Trizol reagent (Sigma). Once obtained, RNAs were subjected to amplification in the presence of CHPV and JEV gene specific-primers by reverse transcriptase-PCR. Primers used to amplify CHPV gene were F (5′-3′) GATCGCGGAGTGGTAGAATATC; R (5′-3′) GAAATCAGCCATGTGTTGTCC, and for JEV GP78, F (5′-3′) TTGACAATCATGGCAAACG; R (5′-3′) CCCAACTTGCGCTGAATAA. Proper negative and positive controls were used and the PCR products were electrophoresed in 2% agarose gel. Purified PCR products (PCR purification kit, Qiagen) of CHPV and JEV positive samples were sent for sequencing. Sequencing was done by Macrogen, Korea.

### Transfection of cells with esiRNA and plasmids

Mouse hnRNPC (EMU173751)-targeting Endonuclease prepared short interfering RNA (esiRNA) and scrambled esiRNA as negative-control were purchased from Sigma-Aldrich. Neuro2A cells upon achieving 60% confluency were undergone transfection with 20 Pico mole of hnRNPC esiRNA along with lipofectamine RNAiMax (Invitrogen) according to manufacturer’s instruction. Following 24 hours of transfection, cells were infected with CHPV and JEV at an MOI of 0.1 and 1 MOI respectively and treated without or with AT. Samples were then processed for immunoblotting analysis after 12 and 24 hours of CHPV and JEV-infection respectively. Viral titer of the cell-culture supernatants collected from these experiments was analyzed by plaque assay. Mock-transfected cells were treated with equivalent volume of Lipofectamine RNAi Max and Opti-MEM in absence of any nucleic acid.

Mouse hnRNPC ORF-harboring expression plasmid, C-Flag tag Plasmid was obtained from Sino Biological (MG51852-CF). E. Coli Top10 strain was transformed with plasmids followed by their amplification and isolation using Plasmid Midi Kit (Qiagen) as per manufacturer’s instructions. 150 ng of plasmids were used for transfection experiments with the aid of Lipofectamine 2000 (Invitrogen, CA, USA) according to manufacturer’s protocol. Empty vector was also transfected and used as negative-control. Post 24 hours of transfection, cells were infected with CHPV and JEV at an MOI of 0.1 and 1 MOI respectively. Virus-infected cells either untreated or AT-treated were incubated for 12 and 24 hours in case of CHPV and JEV infection respectively. The samples were then processed for immunoblotting analysis. Cell-culture supernatants were subjected to plaque assay for evaluating viral titer following incubation with AT for particular time period.

### Statistical analysis

All experiments were performed in triplicate unless otherwise indicated. Student’s two-tailed unpaired t test was performed to assess difference of statistical significance between two groups. Differences amongst multiple groups were examined by one-way ANOVA followed by Holm–Sidak post hoc test. SigmaPlot 11 software was used for data analysis. Data were considered statistically significant when P values were less than 0.05.

## Results

### Atorvastatin conferred protection to animals against viral infections and abrogated inflammatory response

Atorvastatin (AT) treatment was demonstrated to enhance survivability of mice pups following CHPV (Fig. 1A) and JEV infection (Fig. 1B). To assess the protective role of AT, we treated CHPV and JEV-infected mice pups with AT (5 mg/kg of body-weight) administered through intraperitoneal route, 3 hours post-infection (hpi), once daily. Mice pups were carefully monitored for clinical symptoms. AT at a dosage > 5 mg/kg of body-weight resulted in decreased movement and body weight, thus indicating toxicity of AT at higher doses. Data demonstrating the effect of AT treatment at a dosage of 2.5mg/kg of body-weight upon body weight has been provided in Supplementary Fig. 1A and 1B. AT treatment following 4 days of CHPV infection has been observed to enhance survivability by 70 percentage. AT treatment was also found to confer similar protective action against JEV infection. AT when administered for 8 days following infection with JEV, mortality rate plummeted by 80 % when compared to that of only-infected group. The elevated abundance of interleukin-6 (IL-6), monocyte chemoattractant protein-1 (MCP-1), interferon-γ (IFN-γ) and IL-6, MCP-1, IFN-γ and tumor necrosis factor-1 (TNF-α) following CHPV and JEV infection respectively was found to be reversed upon AT treatment (Fig. 1C and 1D) thus demonstrating its immunosuppressive potential in the context of viral infections.

**Fig. 1:**
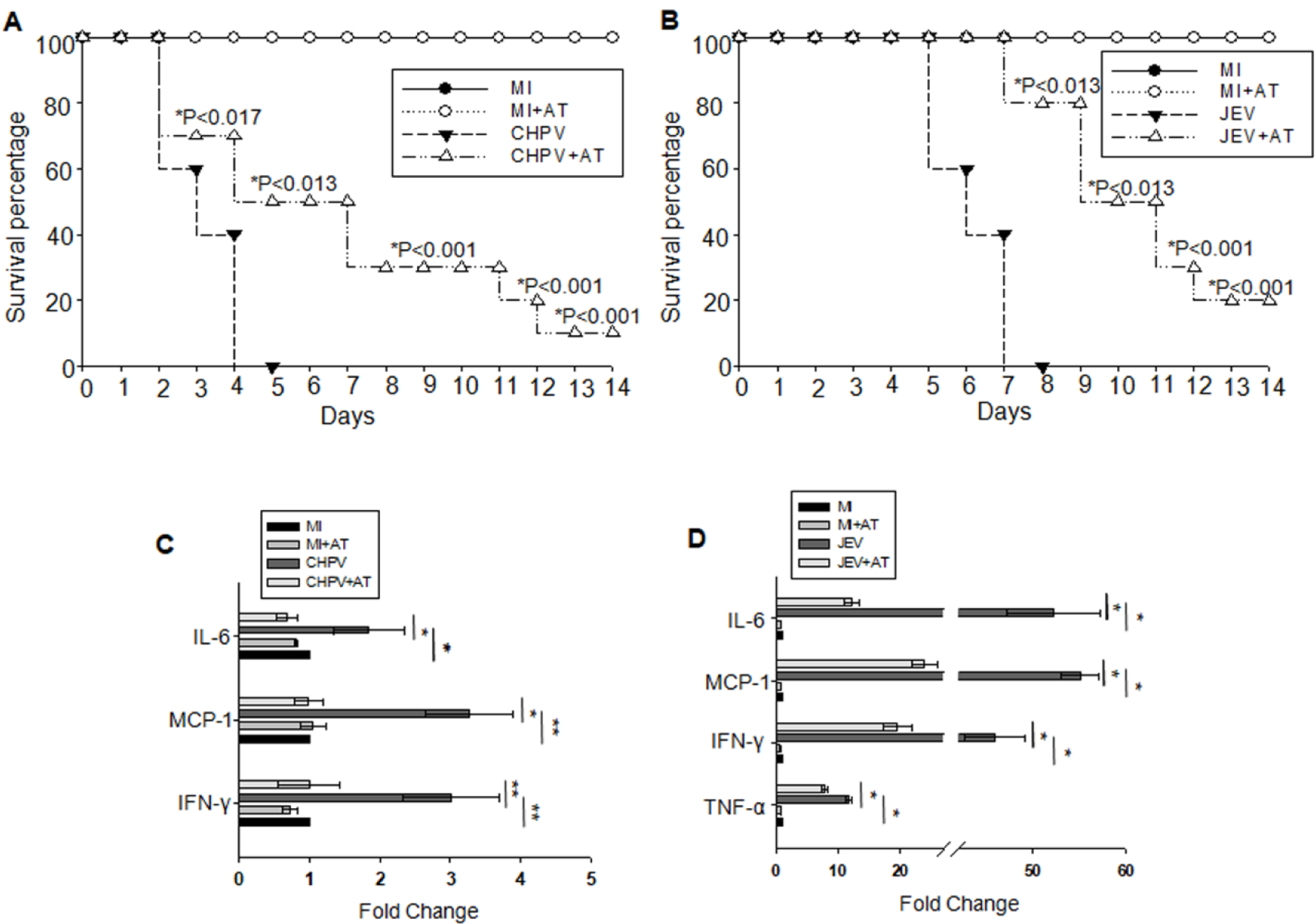
Atorvastatin-treatment increased mice survival and resulted in amelioration of virus-induced neuroinflammation. (A) Mice were infected with CHPV and treated once daily with atorvastatin (AT) at a dosage of 5mg/kg-body weight till day 4 following infection (*p ≤ 0.01), (n=15). (B) JEV-infected mice were administered with AT at a dosage of 5mg/Kg-body weight once daily till day 7 and were monitored for signs of infection (*p ≤ 0.01), (n=15). (C) Brain lysates from CHPV-infected mice exhibiting severe signs (4 dpi), CHPV-infected mice treated with AT, and corresponding control mice were subjected to cytokine bead array and analysed by flow cytometry to observe the status of abundance of IL-6, IFN-γ and MCP-1 (*p ≤ 0.01 and **p ≤ 0.001). Data have been represented as mean ± SD of three independent experiments. (D) Brain lysates from JEV-infected mice demonstrating severe signs of infection (7 dpi), JEV-infected mice treated with AT, and control mice were subjected to cytokine bead array and analysed by flow cytometry to observe the status of expression of IL-6, IFN-γ, MCP-1, TNF-α (*p ≤ 0.001). Data are represented as mean ± SD of three independent experiments. Abbreviations used; Mock Infected (MI), Mock Infected + Atorvastatin (MI + AT), Infected (CHPV or JEV), Atorvastatin Treatment (CHPV + AT or JEV + AT).

### Atorvastatin modulates expression of key stress-related proteins related with stress due to infection severity in vivo

AT treatment resulted in significant reduction of CHPV and JE viral load in the brain as demonstrated by western blot analysis of viral protein abundance, in comparison to the only-infected group. Overexpression of CHPV proteins, viz. glycoprotein (G), nucleoprotein (N) and matrix protein (M), and non-structural protein 3 (NS3) of JEV upon virus infection was found to be reduced by AT administration thus indicating anti-viral role of AT. Consistent with these observations, viral titer measurement by plaque assay analysis demonstrated significantly reduced JEV and CHPV load in the brain of virus-infected animals treated with AT with respect to only-infected animals (Fig. 2C and 2D). AT treatment was also found to reduce virus infection-induced up-regulation of stress-related protein hnRNP C1/C2. Moreover, cellular abundance of Caspase-3 cleaved fragment in virus-infected group was observed to decrease upon AT treatment when compared to that of virus-infected group (Fig. 2A and 2B). Absence of any effect of DMSO upon drug-treated group has been shown in Supplementary Fig. 2A and 2B. Since earlier work by our group demonstrates role of elevation of endoplasmic reticulum resident chaperone GRP78 (Bip) and mitochondrial protein prohibitin in JEV-infected sub ventricular zone (SVZ) of brain (37), we analyzed whether AT treatment reduces cellular death upon viral infection via altering cellular abundance of Bip and prohibitin, and hence ER stress. Albeit both JEV and CHPV infection resulted in increased abundance of Bip and prohibit in virus-infected brain, no change in Bip and prohibitin expression was observed in virus-infected animals upon AT treatment (Supplementary Fig. 2C and 2D). In order to confirm the association of AT treatment’s anti-viral property and its effect upon hnRNPC activity, mice brain samples were subjected to immunohistochemical investigation. Upon JEV and CHPV infection, respective viral proteins were observed to be up -regulated along with cellular abundance of hnRNPC. On the contrary, AT treatment of virus-infected mice culminated into significant reduction of co-localization of viral proteins with hnRNPC (Fig. 3A and 3B).

**Fig. 2:**
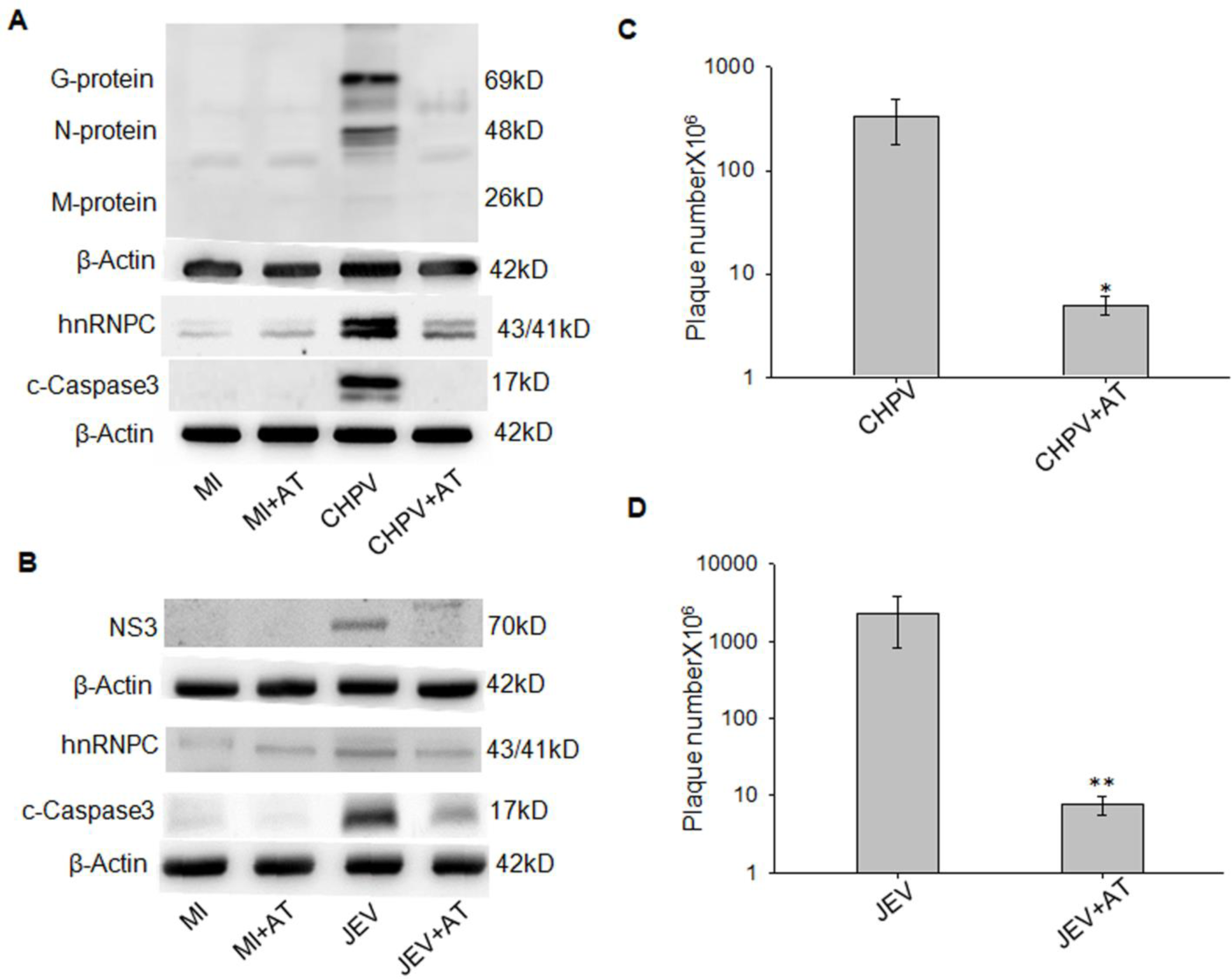
Atorvastatin treatment (once daily at a dosage of 5mg/ kg-body weight) induced reduction in viral (CHPV and JEV) and ER stress-protein expressions *in vivo.* (A) and (B) Mice were divided into 4 groups, viz. Mock-infected (MI), mock-infected but treated with AT (MI+AT), CHPV/JEV-infected and CHPV/JEV-infected mouse treated with AT (AT treatment). Following 3 hours of infection with CHPV/JEV, AT was administered once daily. Signs appeared following 3 and 7 days of CHPV and JEV infection, respectively. Following mice brain collection, they were subjected to immunoblotting for analysis of effect of AT upon cellular apoptosis, ER stress and viral propagation. Viral propagation was assessed by estimating glycoprotein (G), nucleoprotein (N), Matrix protein (M) abundance in case of CHPV and NS3 for JEV. Apoptosis and ER stress was analysed by evaluating the abundance of cleaved-caspase-3 and hnRNPC1/2. (C) and (D) demonstrate the effect of AT treatment upon virus production in the mouse brain. Plaque assay was performed using supernatant collected from brain homogenate of CHPV/JEV-infected and AT-treatment group. (*p ≤ 0.001 and **p ≤ 0.026). Data represented here as mean ± SD of three individual experiments.

**Fig. 3:**
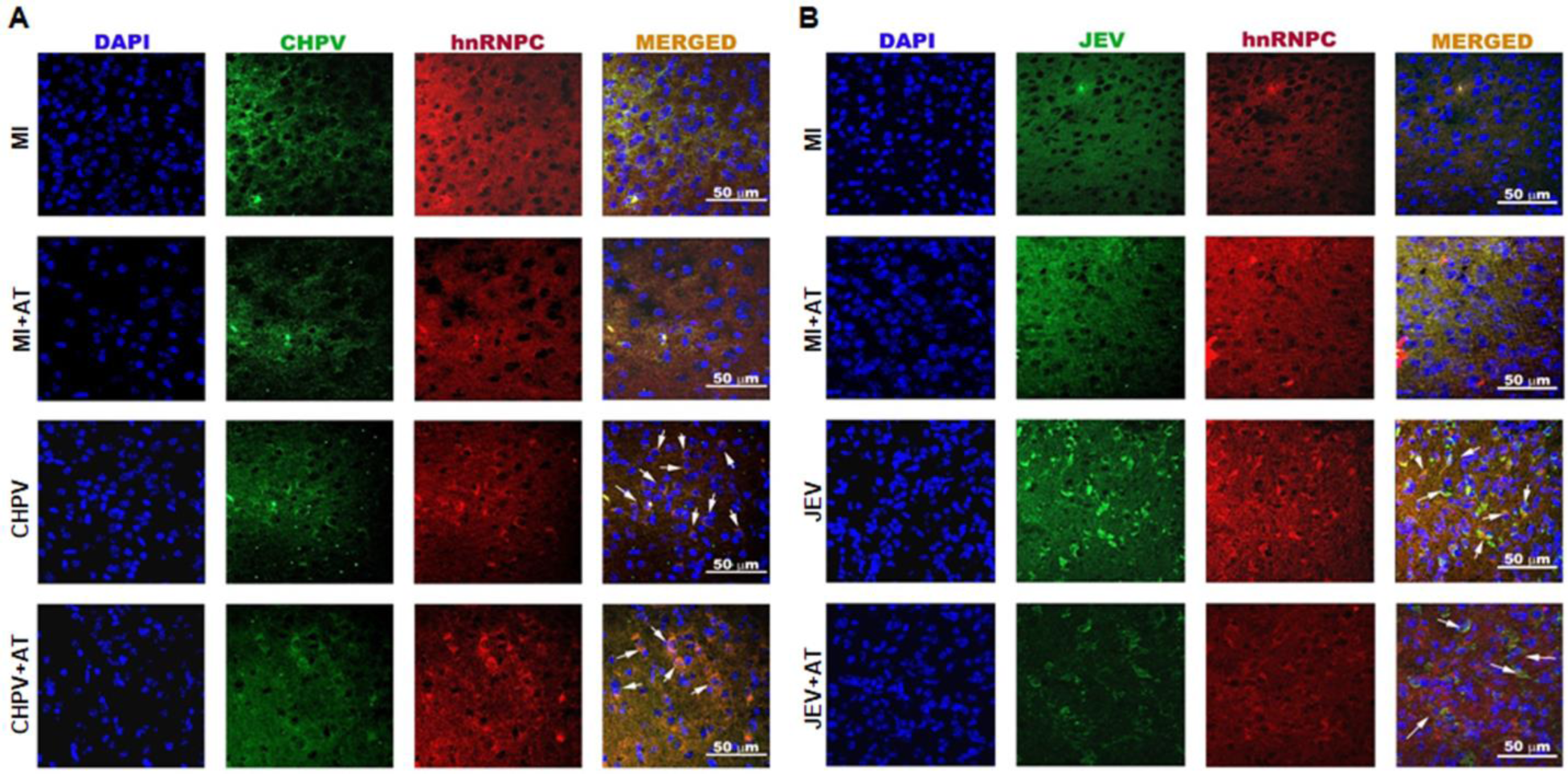
Reduction in hnRNPC abundance upon treatment with AT in virus-infected brain. (A and B) Balb C mice were infected with CHPV and JEV. AT treatment was administered once daily. Following appearance of signs of virus infection, brain samples were collected followed by fixation, generation of brain sections, permeabilization and stained with antibodies for detection of viral proteins and hnRNPC Scale bar 50 µm. Data presented are representative of three independent experiments.

### Atorvastatin ameliorates expression of key protein related with stress and viral load in vitro

Determination of non-toxic dosage of AT was determined by performing dose-response experiment. Analysis of dose-response curve demonstrating toxicity imparted on Neuro2A cell line by various concentrations of drug showed that AT concentration greater than 2µM resulted in decrease in cell viability. Whereas, AT dosage at concentration less than or equal to 2µM was found not to reduce cellular viability (supplementary fig. 3A). We further examined the effect of DMSO upon cellular biology in response to infection, and found that DMSO does not alter CHPV and JEV propagation and infection-induced cell death *in vitro* (Supplementary Fig. 3B and 3C). Treatment with AT (2 µM) reduced generation of intracellular reactive oxygen species when compared to that in infected groups (JEV and CHPV infection) (Supplementary Fig. 4A and 4B). AT treatment was found to result in reduction of CHPV and JEV protein abundance in a dose-dependent fashion as shown by the immunoblotting analysis of neuro2a infected and drug-treated (1 µM and 2 µM) cells at 12 and 24 hours post-infection. Expression of hnRNP C1/C2 protein in AT-treated cells was found to decrease with respect to virus-infected cells. Consistent with our *in vivo* data, AT treatment culminated into reduced activation of caspase-3 when compared to that in virus-infected cell as shown by reduced abundance of cleaved-caspase-3 fragment (Fig. 4A and 4B). AT treatment also helped reduce cellular viral propagation as shown by immunoblotting analysis of glycoprotein (G), nucleoprotein (N) and matrix proteins(M) of CHPV and non-structural protein 3 (NS3) of JEV. The aforementioned observations were also supported by plaque assay analysis thus concluding the anti-viral action of AT upon CHPV and JEV infection (Fig. 4C and 4D). Surprisingly, we also found that elevated level of hnRNP C1/C2 and cleaved caspase-3 due to thapsigargin-induced ER stress was diminished by the treatment of atorvastatin (2 µM), *in vitro* (Fig: 4E).

**Fig. 4:**
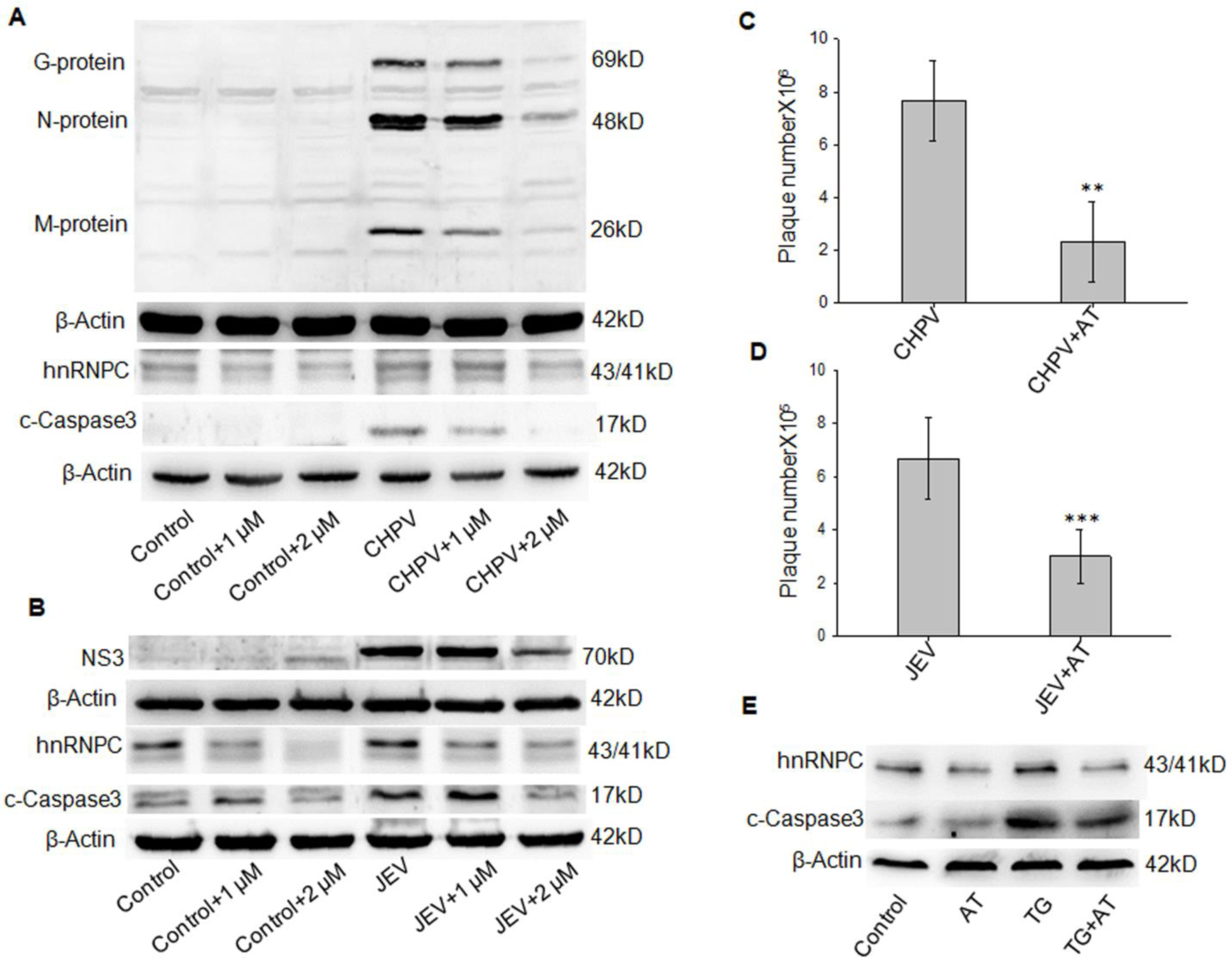
Atorvastatin induced reduction in viral (CHPV and JEV) and ER stress-protein expressions *in vitro.* (A) and (B) Six groups of Neuro2A cells were studied; Control (CT), Control-cells treated with AT (1 µM and 2 µM), virus-infected cells (CHPV/JEV infection) and virus-infected cells treated with AT (1 µM and 2 µM). Cells were incubated with viruses for 2 hours followed by their removal. Cells were then maintained in maintenance media containing indicated concentrations of AT. CHPV and JEV-infected cell samples were collected 12 and 24 hours post infection respectively. Following sample collection, they were subjected to immunoblotting for analysis of effect of AT upon cellular apoptosis, ER stress and viral propagation. Viral propagation was assessed by estimating glycoprotein (G), nucleoprotein (N), Matrix protein (M) abundance in case of CHPV and NS3 for JEV. Apoptosis and ER stress were analysed by evaluating the abundance of cleaved-caspase-3 and hnRNPC1/2. (C) and (D) Effect of AT treatment upon viral propagation was also studied using plaque assay analysis of the cell culture supernatants of the only-infected and AT-treated infected samples. Concentration of AT used for treating infected-neuro2A cells was 2 µM. (*p ≤ 0.020 and **p ≤ 0.038). Data are represented as mean ± SD of three individual experiments. (E) Effect of thapsigargin (TG) treatment upon ER stress and apoptosis in absence as well as presence of AT was studied using immunoblotting analysis. Cells were pre-treated with TG (1 µM) followed by incubation in maintenance media containing/lacking 2 µM AT for 24 hours. Status of ER stress and apoptosis were evaluated by assessing abundance of hnRNPC and cleaved-Caspase 3. β-actin served as loading control.

### Atorvastatin abrogates virus infection in primary cortical neurons

To further validate our findings, we evaluated the association of virus infection with hnRNPC1/C2 up regulation in primary culture of mouse cortical-neurons. Mouse cortical-neuronal cells were isolated and cultured. Cells were then left either uninfected or infected with JEV or CHPV for 24 and 12 hours respectively. Uninfected cells were also treated with AT, which served as a negative-control. Separate groups were also included in the experiment to assess the effect of AT treatment upon virus infections. Cell samples were then subjected to immunocytochemical analysis using anti-hnRNPC anti-CHPV and anti-JEV antibodies. As hypothesized, enhanced hnRNPC abundance exhibited by virus infected-cells was reversed by AT treatment (Fig. 5A and 5B).

**Fig. 5:**
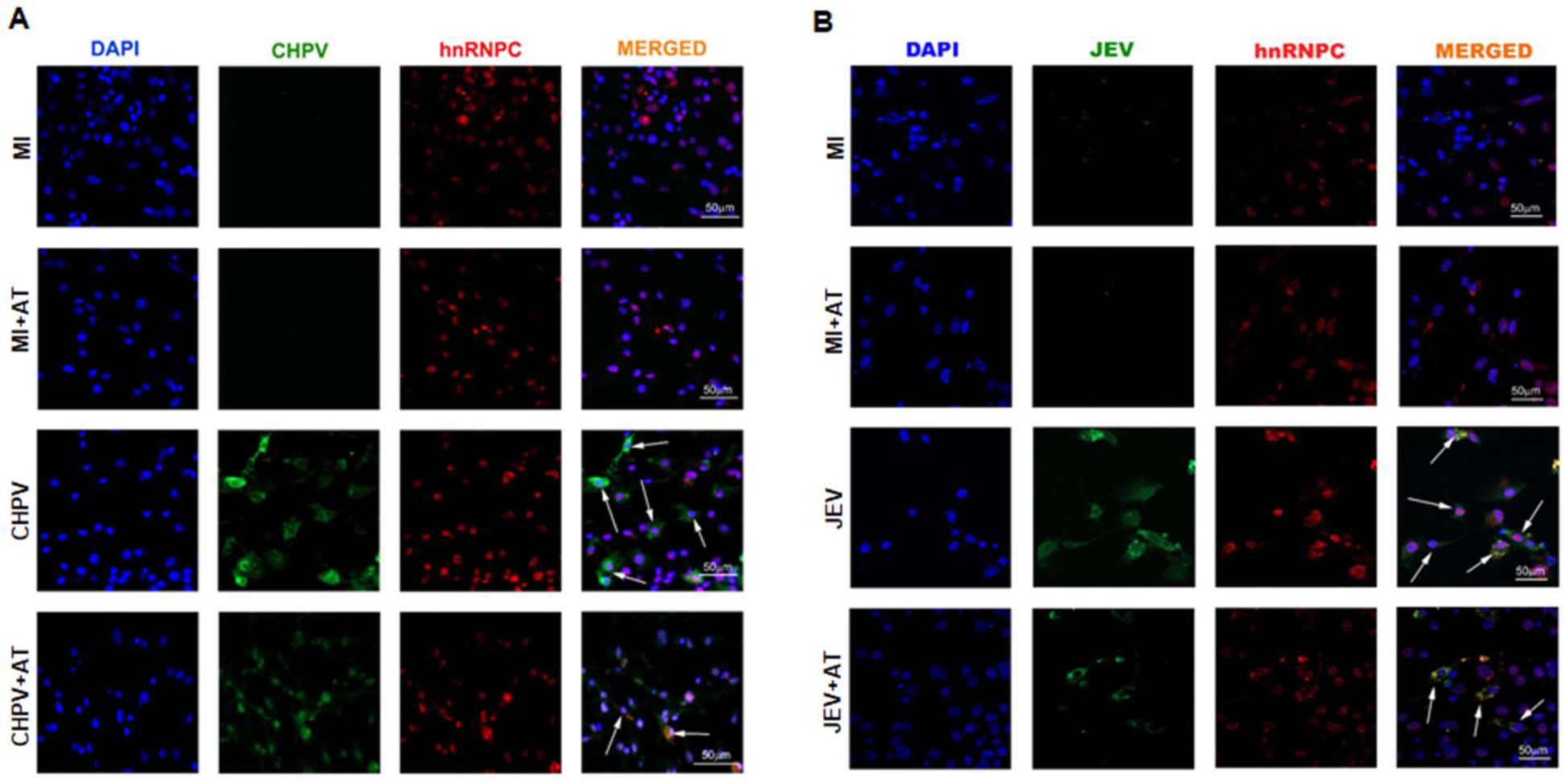
AT treatment abrogated cellular hnRNPC abundance and viral propagation in primary culture of mouse cortical neurons. (A) and (B). Cortex from 2-days old mice pups were excised followed by isolation of cortical cells. Cortical cells were then cultured on PDL-coated petri dishes. Cells were then infected with CHPV (A) and JEV (B) for 12 and 24 hours respectively. AT-treated cells were maintained in AT-containing maintenance media following incubation with virus particles. Cells were then collected and subjected to immunocytochemical analysis for viral proteins and hnRNPC. Scale bar used: 50 µm. Data presented are representative of three independent experiments.

### Atorvastatin negatively regulates CHPV and JEV replication in an hnRNPC1/2-associated fashion, in vivo and in vitro

In order to assess the effect of AT treatment upon hnRNPC1/2 and viral genomic RNA interaction, RNA co-immunoprecipitation (RNA-co-IP) was performed using antibody against hnRNPC1/2. *In vivo* samples were subjected to undergo RNA-co-IP (Fig. 6A and 6B). Neuro2A cells were infected with JEV and CHPV for 24 and 12 hours respectively. Infected cells were then either treated with AT or left untreated. Following the above mentioned steps, hnRNPC1/2 was pulled down using anti-hnRNPC antibody and its interaction with viral genomic RNA was analyzed (Fig. 7A and 7B). Use of control IgG and absence of reverse transcription reaction in RNA-co-IP experiment resulted in no signal detection, and were used as negative-controls. Sequencing the specific bands following gel extraction revealed sequences possessing 89% and 99% sequence similarities with CHPV and JEV genomic sequence respectively, *in vivo* (Fig. 6A and 6B). Sequence similarities of 91% and 99% with CHPV and JEV genome respectively were observed after bands were subjected to sequencing following RNA-co-IP experiments using CHPV and JEV respectively, *in vitro* (Fig. 7A and 7B). Consistent with our hypothesis, both *in vivo* and *in vitro* experiments demonstrated that AT treatment resulted in diminution of genomic RNA abundance obtained following pulling down hnRNPC protein in comparison to untreated but infected group, thus pointing towards the fact that AT treatment might affect viral propagation by interfering with the formation of hnRNPC-possessing viral genome replication complex.

**Fig.6.**
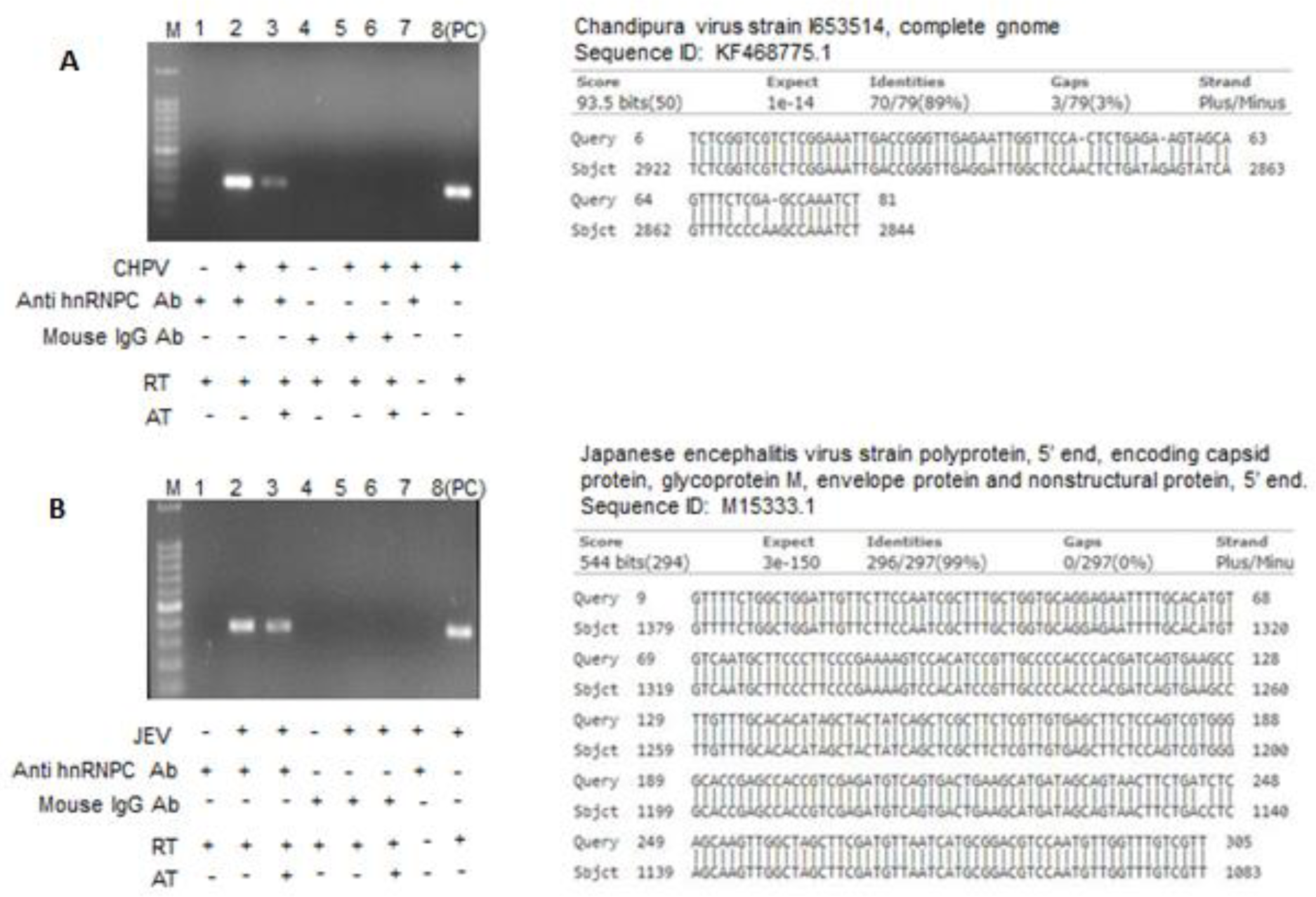
AT treatment resulted in reduction of hnRNPC-JEV/CHPV genome interaction *in vivo*. (A) Mouse brain samples from mock-infected, CHPV-infected and infected-mice treated with AT were collected 4 days after infection followed by homogenization, crosslinking of RNA and proteins, release of the cross linked complex by sonication and immunoprecipitation of hnRNPC bound to its interacting partners. RNA was then isolated from the hnRNPC-containing solution and was subjected to reverse transcription followed by amplification using viral-genome specific primers. Band (∼296 bp) obtained upon amplification using CHPV-specific primer from infected-mouse brain was used as a positive control (PC). Size of the bands obtained upon immunoprecipitation of hnRNPC was found to be identical to PC band thus indicating detection of CHPV genome. Sequencing data of the identified band exhibited more than 89% similarity with CHPV strain I653514 genome. (B) Mouse brain samples from mock-infected, JEV-infected and infected-mice treated with AT were collected 7 days after infection followed by processing similarly to that stated for (A). Band (∼400 bp) obtained upon amplification using JEV-specific primer from infected-mouse brain was used as a positive control (PC). Size of the bands obtained upon immunoprecipitation of hnRNPC was found to be identical to PC band thus indicating detection of JEV genome. Sequencing data of the identified band displayed more than 99% similarity with JEV strain polyprotein 5’ end.

**Fig.7.**
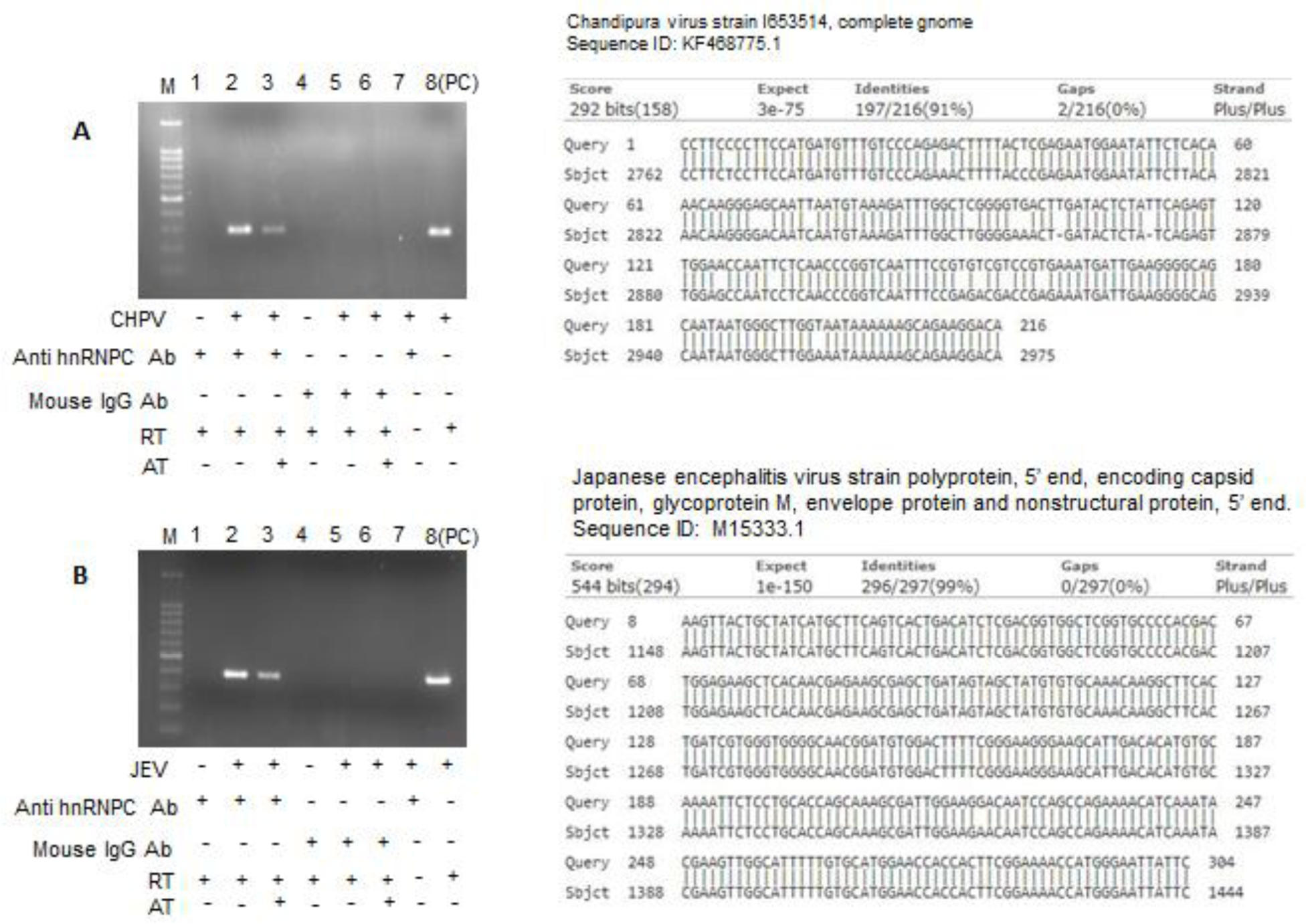
AT treatment *in vitro* reduced CHPV and JEV genomic RNA interaction with hnRNPC. (A) CHPV-infected Neuro2A cells were collected following 12 hours of infection followed by RNA-protein crosslinking, cell lysis and immunoprecipitation of hnRNPC using anti-hnRNPC antibody. RNA was isolated from hnRNPC-containing complex followed by amplification using CHPV-specific primers. CHPV-infected cells were subjected to reverse transcription-polymerase chain reaction using CHPV-specific primers, and the band obtained was used as positive control (PC). Bands obtained upon immunoprecipitation using anti-hnRNPC antibody exhibited size (∼296 bp) similar to that of (PC) band. Sequencing data of the identified band revealed more than 91% similarity to that of CHPV strain I653514 genome. (B) JEV-infected Neuro2A cells collected following 24 hours of infection were processed as stated in (A). Amplification using JEV genome-specific primers resulted in appearance of bands having size similar ((∼400 bp) to that of (PC) band for JEV genome fragment. Sequencing data of the identified band revealed more than 99% similarity to that of JEV strain polyprotein 5’ end.

### Validation of function of hnRNP C1/C2 in JEV and CHPV viral infection manifestation and its modulation by atorvastatin

Since AT has been observed to reduce expression of ER stress-generated hnRNPC1/2 in the context of virus infections, we used hnRNPC1/2 knock-down and over-expression studies to decipher mechanistic way of AT action upon viral propagation. Immunoblotting analysis exhibited decrease in viral proteins (G, N and M proteins of CHPV and NS3 of JEV) abundance upon hnRNP C1/C2 knockdown in Neuro2A cells. Further decrease in viral proteins was observed in samples where concomitant knockdown of hnRNPC by siRNA delivery was performed with AT treatment (2 µM) (Fig. 8A and 8B). These findings were further supported by virus titer analysis where CHPV (at 12hpi) and JEV (at 24hpi) propagation was found to decline upon hnRNPC knockdown with or without AT treatment *in vitro* (Fig. 8C and 8D). Overexpression of hnRNPC1/2 was performed in order to evaluate the effect of hnRNPC1/2 upon viral propagation. Plasmid possessing hnRNPC1/2 construct was used for transfection of neuro2A cells. Immunoblotting using antibodies against G, N and M proteins of CHPV and NS3 of JEV revealed positive role played by hnRNPC overexpression in promoting viral multiplication, even in the face of ATR treatment (2 µM) (Fig. 9A and 9B). Consistent with the immunoblotting experiments, plaque assay analysis also demonstrated enhanced CHPV (12 hpi) and JEV (24 hpi) reproduction upon hnRNPC1/2 overexpression irrespective of the presence of atorvastatin *in vitro* (Fig. 9C and 9D).

**Fig. 8:**
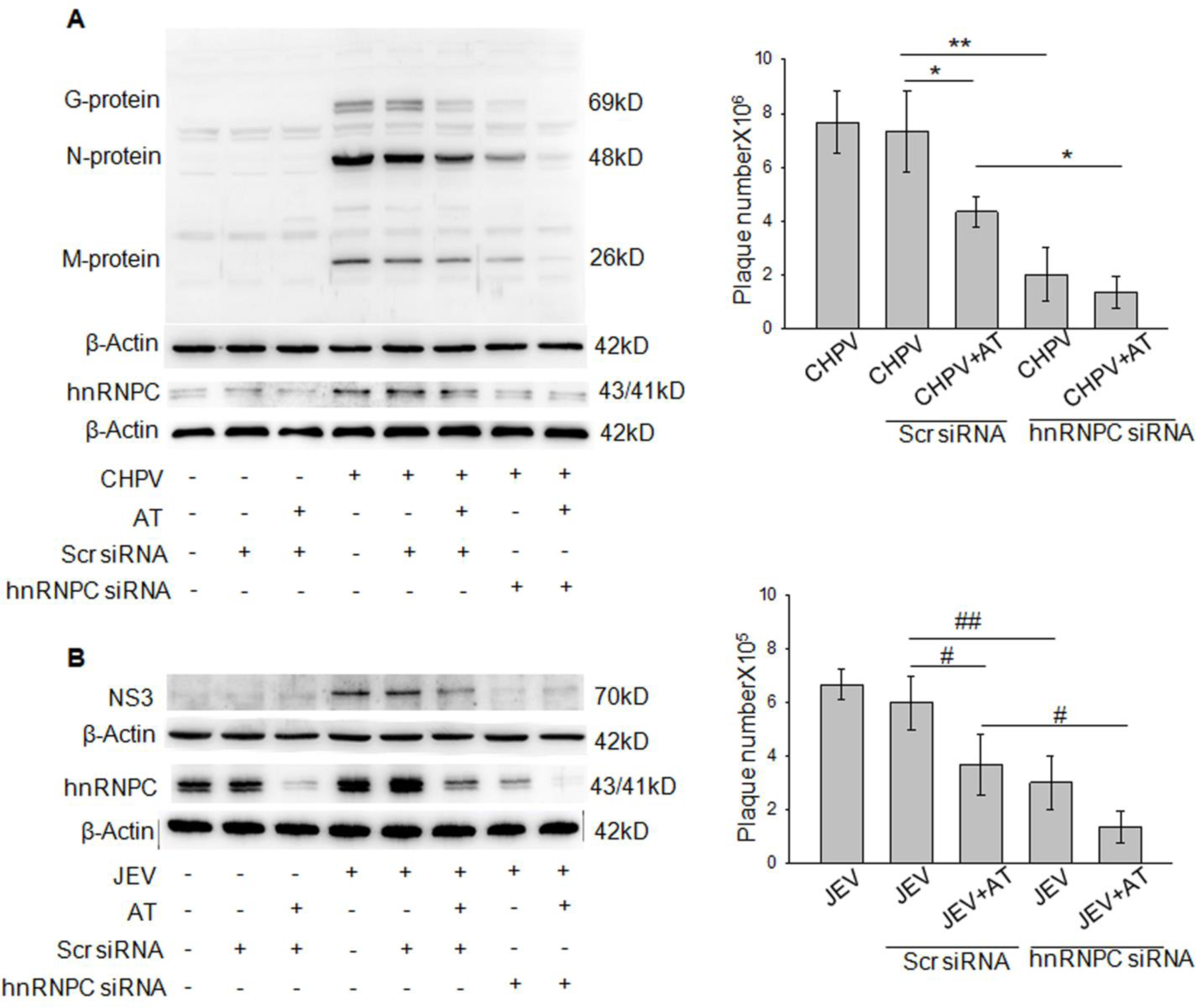
AT-induced decrease in viral (CHPV and JEV) propagation is mediated by down regulation of host hnRNPC1/2 *in vitro*. (A). esiRNA treatment of Neuro2A cells for 24 hours was performed prior to CHPV infection (0.1 MOI) for 12 hours. Amount of esiRNA used was 20 Pico moles. Followed by virus infection along with respective treatments, cells were collected and subjected to immunoblotting technique for evaluation of effect of AT treatment upon CHPV propagation. β-actin served as loading control. (B) Neuro2A cells were transfected with esiRNA similarly to that in (A) and infected with JEV (1 MOI) for 24 hours. Following infection, cells were harvested and processed for immunoblotting analysis. Antibody against NS3 protein was used for determination of level of JEV propagation upon knock-down of hnRNPC. β-actin served as loading control. (C) Neuro2A cells were transfected with esiRNA and either left uninfected or infected by CHPV (C) and JEV (D) in presence or absence of AT (2 µM). Following 12 and 24 hours of CHPV (0.1 MOI) and JEV (1 MOI) infection respectively, cell supernatant from different samples were collected and analysed for viral titer by plaque assay. (*p ≤ 0.03, **p ≤ 0.001, ^#^p ≤ 0.04, ^##^p ≤ 0.01). Data represented mean ± SD of three independent experiments.

**Fig. 9:**
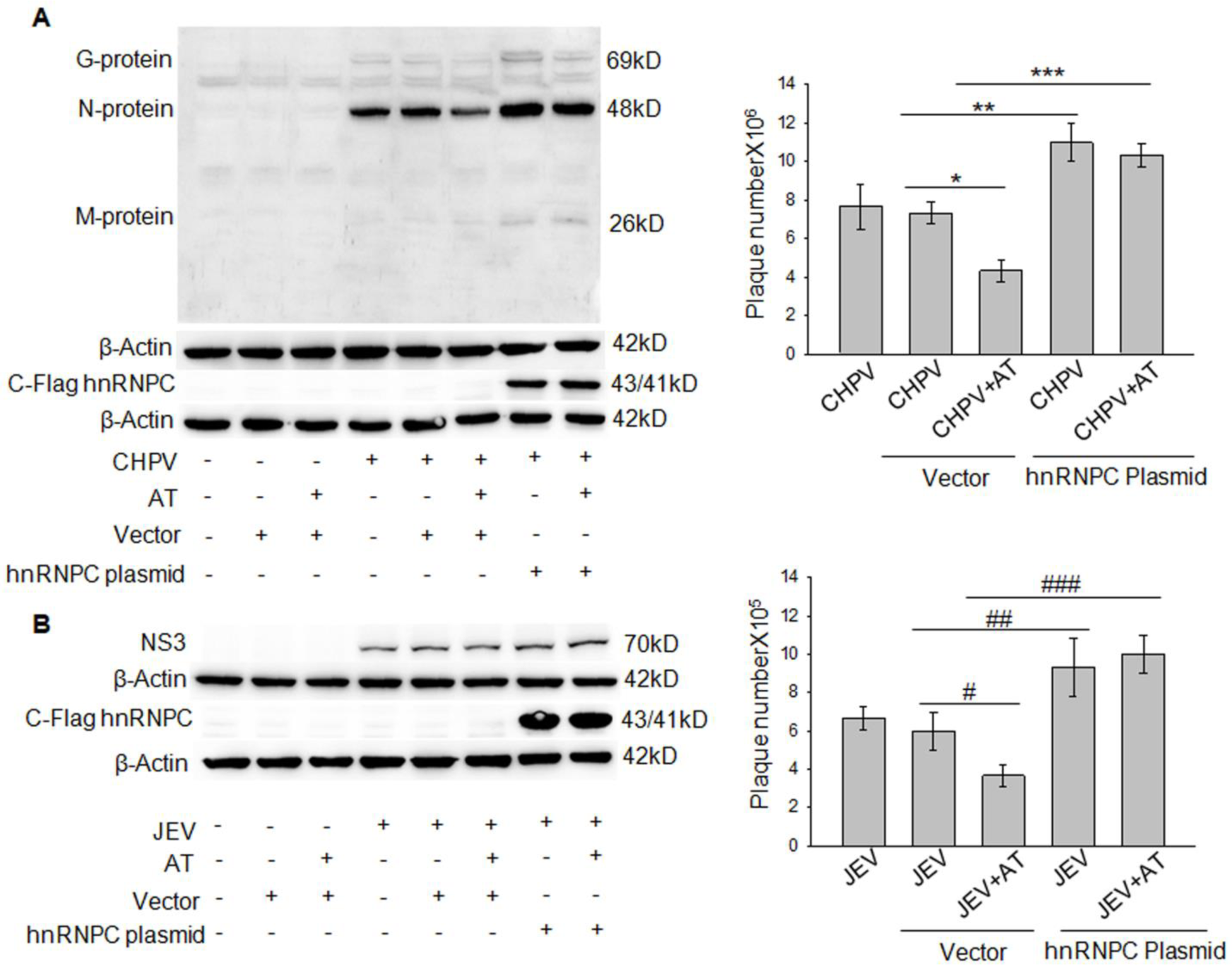
Overexpression of hnRNPC abrogated effect of AT treatment upon virus propagation *in vitro*. Neuro2A cells were transfected with plasmids possessing c-flag-tagged hnRNPC gene (150ng) for 24 hours. Following transfection, cells were infected with CHPV (A) at 0.1 MOI and JEV (B) at 1 MOI for 12 and 24 hours respectively. Cells were then collected and subjected to immunoblotting analysis for assessment of effect of AT (2 µM) upon viral propagation in the context of altered hnRNPC expression. Antibodies against G, N and M proteins were used for detection of CHPV protein abundance. Anti-NS3 antibody was utilized for detection of JEV protein load. Anti-flag-c antibody was used to determine overexpression of hnRNPC upon plasmid transfection. β-actin served as loading control. (C) and (D) Neuro2A cells were transfected with hnRNPC-expressing plasmid and either left uninfected or infected by CHPV (C) and JEV (D) in presence or absence of AT(2 µM). Following 12 and 24 hours of CHPV (0.1 MOI) and JEV (1 MOI) infection respectively, cell supernatant from different samples were collected and analysed for viral titer by plaque assay. Values represented here as mean ± SD of three independent experiments.

### Atorvastatin acts to reduce virus-induced miR-21 expression in an hnRNPC-dependent fashion

In addition to acting as a positive regulator of various positive-stranded RNA virus replication (38), hnRNPC is also reported to modulate expression of a battery of other host molecules (39, 40) thus underscoring its importance in alteration of various cellular processes. Report exists (38) demonstrating the role of hnRNPC in mediating downregulation of PDCD4 protein in a miR-21-dependent mechanism. On the other hand, reduction of PDCD4 has been documented to promote apoptosis as a result of translation of procaspase-3 (41). In order to decipher the mechanistic details regarding mode of AT action, effect of AT treatment upon miR-21 abundance was studied both *in vivo* and *in vitro* (Fig: 10A and 10B), in the context of CHPV and JEV infections. Balb C mice were treated with PBS (MI), PBS and atorvastatin (MI+AT), CHPV or JEV (Fig. 10A & 10B), CHPV/JEV and atorvastatin (CHPV/JEV + AT). Both virus and atorvastatin treatment were administered via intra-peritoneal route. AT treatment was administered once daily starting from 3 hours post infection. Following 3 and 7 days of CHPV and JEV infection respectively, brain samples were harvested and subjected to total RNA isolation. RNA isolated was subjected to reverse transcription reaction followed by qRT-PCR for analysis of miR-21 abundance. Infection with both CHPV and JEV resulted in increased abundance of miR-21 in comparison to virus-infected animals receiving AT treatment (Fig: 10A). Consistent with *in vivo* data, *in vitro* experiments using neuro2A cell line also revealed similar role of AT treatment upon CHPV/JEV infection-elicited miR-21 upregulation (Fig: 10B). Analysis of human autopsy tissue samples also confirms upregulation of miR-21 upon JEV infection (Fig: 10C). In order to confirm the role of hnRNPC in elevating infection-induced miR-21 expression, neuro2A cells were either mock-transfected or transfected with hnRNPC-specific/scrambled esiRNA followed by infection with CHPV or JEV. While virus-infected samples exhibited concomitant upregulation of miR-21 and hnRNPC, AT treatment was observed to reduce expression of both in the face of CHPV (Fig: 10D) and JEV infections (Fig: 10E). In addition to that, downregulation of host protein hnRNPC in the presence of AT treatment resulted in further decrease in miR-21 expression in case of CHPV (Fig: 10D) and JEV infection (Fig: 10E) with respect to cells transfected with scrambled esiRNA. To revalidate the role played by hnRNPC in triggering miR-21 upregulation upon CHPV and JEV infections, neuro2A cells were either mock-transfected or transfected with empty/hnRNPC-coding plasmids. Upregulation of hnRNPC abundance in neuro2A cells resulted in restoration of miR-21 upregulation upon CHPV (Fig: 10F) and JEV (Fig: 10G) infections despite of atorvastatin treatment. On the other hand, cells on being transfected with empty plasmid were seen unable to maintain virus-induced elevation of miR-21 abundance thus pointing towards role of hnRNPC in regulating miR-21 expression in a positive manner.

**Fig 10:**
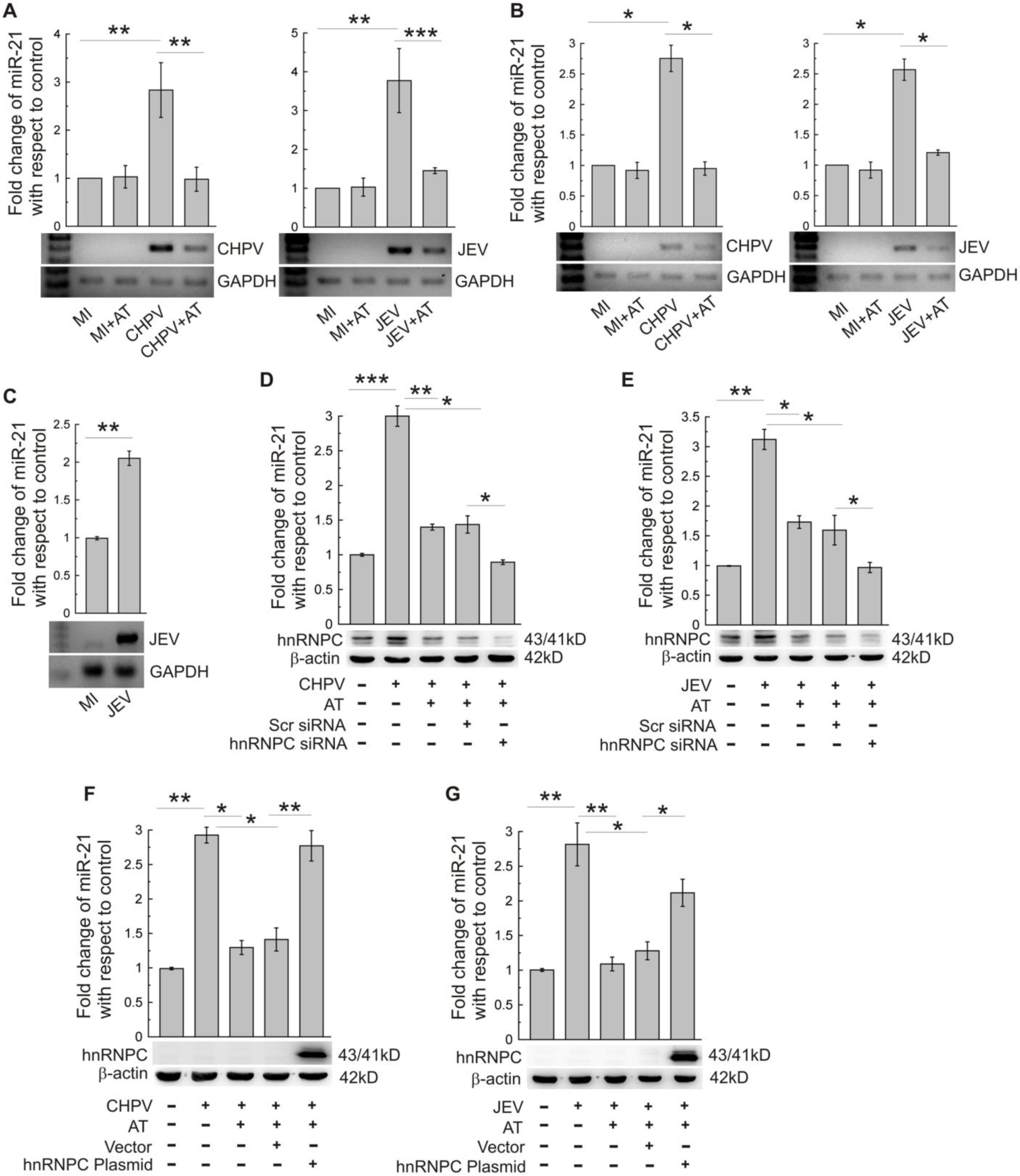
Atorvastatin suppresses virus-induced miR-21 expression through hnRNPC. (A) BALB/c mice were treated with PBS (MI), PBS with atorvastatin (MI+AT), infected with CHPV (CHPV) and CHPV-infected along with atorvastatin treatment (CHPV+AT). Following 3 days post infection, brain samples were collected and miR-21 abundance was analyzed using qRT-PCR. In another similar set of experiments, infection was performed by JEV followed by collection of brain samples at day-7 after infection and miR-21 expression was analyzed by qRT-PCR. In order to assess level of viral propagation, presence of CHPV or JEV RNA was determined by RT-PCR (lower panels). GAPDH expression was verified as loading control. \*\**P*< 0.01, \*\*\**P* < 0.001. (B) N2A cells were left untreated or treated with atorvastatin (2 µM) following incubation of cells with JEV/CHPV for 2. Following 12 and 24 hours of infection with CHPV and JEV respectively, cell samples were subjected to qRT-PCR analysis to determine miR-21 abundance. The degree of viral load was evaluated by RT-PCR of viral RNA (lower panels). GAPDH was used as internal control. \*\**P*< 0.05. (C) miRNA was isolated from uninfected (MI) and JEV-infected human brain sections and miR-21 expression was determined by qRT-PCR. Data are representative of two different brains per group. \**P* < 0.05 (Student’s *t* test) compared to uninfected human brain. RT-PCR of viral RNA was performed to detect presence of viral infection. GAPDH was used as loading control. (D and E) N2A cells were either left untransfected or were transfected with scrambled (Scr) siRNA/esiRNA specific for hnRNPC before being infected with CHPV (D) and JEV (E) followed by atorvastatin treatment. After 12 and 24 hours of infection with CHPV and JEV respectively, miR-21 expression was determined by qRT-PCR analysis. In order to confirm whether effective transfection has taken place, cellular proteins were isolated and subjected to immunoblotting analysis (lower panels) with anti-hnRNPC\**P*<0.05, \*\**P*< 0.01, \*\*\**P* < 0.001. β-actin served as a loading control. (F and G) Cells were either left untransfected or transfected with indicated plasmids. 24 hours following transfection, cells were left uninfected or infected with CHPV (F) and JEV (G) followed by treatment of atorvastatin. Preceded by 12 and 24 hours of infection by CHPV and JEV respectively, miR-21 expression was analyzed by qRT-PCR. Immunoblotting analysis was also performed to confirm effective transfection of respective plasmids. β-Actin used as a loading control.\**P*<0.05, \*\**P*< 0.01.Data represented here as mean ± SD of three individual experiments.

### Atorvastatin treatment culminates into gain in PDCD4 expression and decreased Caspase-3 activation as a result of miR-21 downregulation

*In vivo* data obtained from Balb C mice infected with CHPV and JEV displayed decrease in PDCD4 abundance at the protein level upon virus infections (Fig: 11A). However, RNA level of PDCD4 did not exhibit a similar decline in abundance upon CHPV and JEV infections. Our *in vivo* data was further corroborated by *in vitro* experiments demonstrating decline in PDCD4 abundance at the protein level upon CHPV and JEV infections (Fig: 11B). To analyze the effect of miR-21 downregulation upon PDCD4-mediated cell death in the context of CHPV and JEV infections, neuro2A cells were either left uninfected, infected, infected along with transfection with miR-21 inhibitor/control-inhibitor. Interestingly, inhibition of miR-21 in the face of CHPV infection resulted in restoration of PDCD4 back to its normal level and reduced Caspase-3 activation with respect to CHPV-infected samples (Fig: 11C). Whereas, transfection of CHPV-infected cells with inhibitor-control resulted in no significant changes in PDCD4 protein abundance or Caspase-3 activation status (Fig: 11C). Similar effect of miR-21 knockdown upon PDCD4 expression and Caspase-3-mediated cell death was observed upon infection with JEV (Fig: 11D). Expression status of miR-21 was checked in order to confirm efficiency of anti-miR transfection (Fig: 11C and 11D). CHPV and JEV-infected Neuro2A cells treated with atorvastatin were found to prevent reduction in PDCD4 abundance when compared to that in cells infected CHPV and JEV in the absence of atorvastatin treatment both *in vitro* and *in vivo* (Fig: 11E and 11F). Moreover, treatment with atorvastatin was discovered to abrogate the increase in miR-21 expression upon CHPV and JEV infections, both *in vitro* and *in vivo* (Fig: 11E and 11F).

**Fig. 11:**
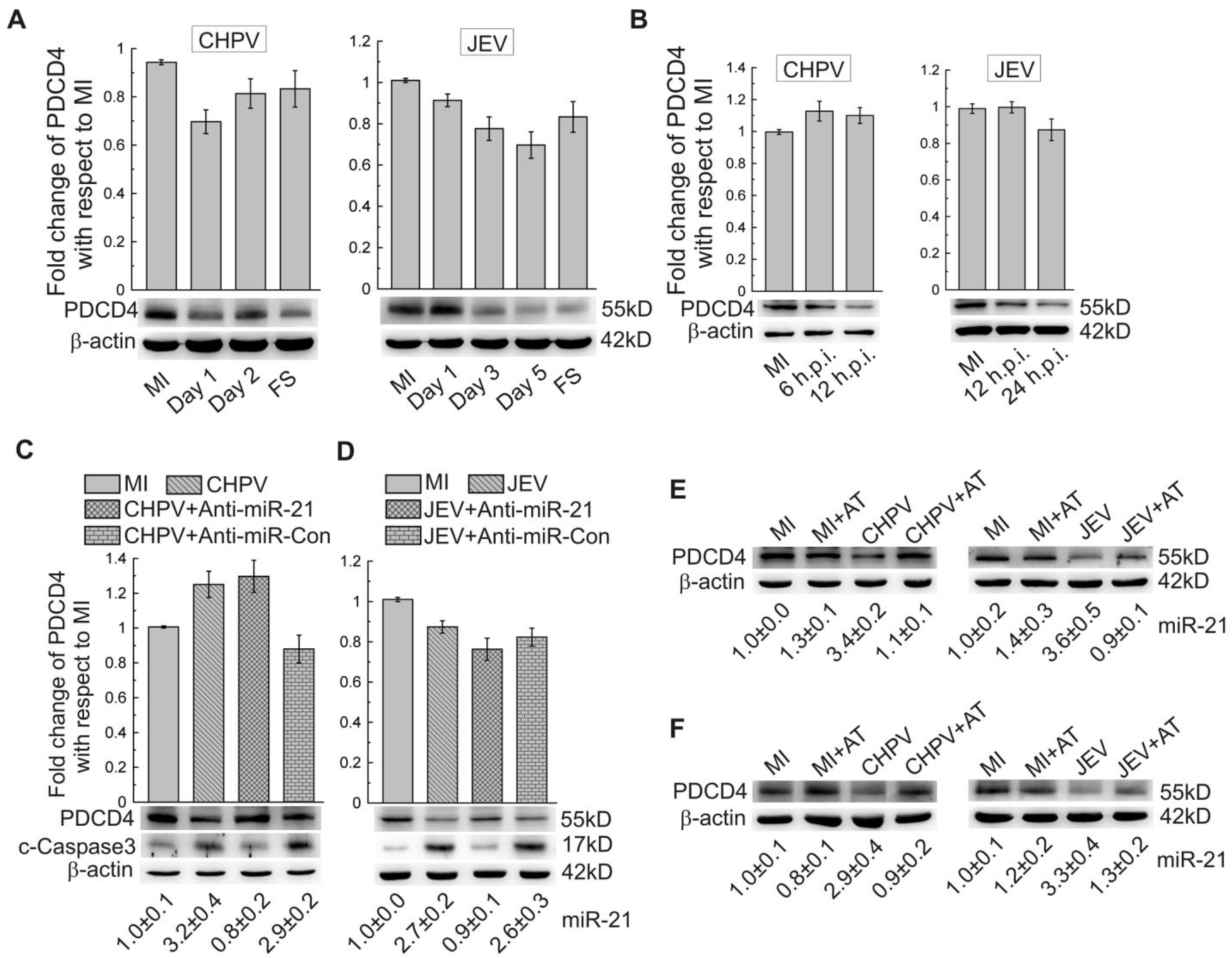
Atorvastatin reduces miR-21 mediated PDCD4 loss and decreases caspase-3 activation. (A) Mice were left uninfected (MI) or were infected with CHPV or JEV. Mice brains were harvested following indicated time periods of infection and subjected qRT-PCR and western blot to determine the expression of PDCD4 mRNA and protein (lower panels) respectively. β-actin served as loading control. FS: full-symptoms. (B) N2A cells were either mock-uninfected (MI) or infected with CHPV or JEV for indicated time periods. Top: PDCD4 mRNA abundance was measured by qRT-PCR analysis. Bottom: Proteins isolated were subjected to immunoblotting procedure for evaluation of expression of PDCD4. β-actin was used as loading control. (C and D) N2A cells were transfected either with miR-21 (Anti-miR-21) or the inhibitor-control (Anti-miR-Con) and then were infected with CHPV (C) and JEV (D) for 12 hours and 24 hours respectively. The cellular RNA was analyzed by qRT-PCR to determine the abundances of PDCD4. Western blotting (lower panels) was performed to evaluate expression of PDCD4 and cleaved caspase3 (c-Caspase3). β-actin was used as a loading control. The expression of miR-21 in each set of experiments was analyzed by qRT-PCR and is shown below the blots to confirm effective transfection. (E and F) N2A cells (E) were mock-uninfected (MI) or infected with CHPV and JEV. Followed by 2 hours incubation with virus, cells were washed thoroughly and left untreated or were treated with atorvastatin (2 µM). Following 12 and 24 hours of infection with CHPV and JEV respectively, proteins isolated from cells were analyzed by western blotting to detect PDCD4 expression. β-actin was served as a loading control. The expression of miR-21 as determined by qRT-PCR for each set of cells is shown below the blots. Mice were either mock-infected (MI) with PBS or infected with CHPV (CHPV) and JEV (JEV) or CHPV and JEV-infected along with atorvastatin treatment (CHPV+AT/JEV+AT) (F). Brain samples were collected after 3 days and 7 days of CHPV and JEV infection respectively. Immunoblot was performed to analyze PDCD4 protein expression as well as miR-21 expression analyzed by qRT-PCR in respective samples was shown below the blots. β-actin was used as a loading control. Data are means ± SD of three independent experiments.

## Discussion

Earlier reports have repeatedly underscored the role of atorvastatin (AT) in amelioration of different pathological states in a fashion independent of its lipid lowering actions. Since AT has been reported to cross the blood brain barrier *in vivo*, repurposing of the same in the setting of traumatic brain injury (TBI) has resulted in enhanced neurogenesis thus leading to decreased injury-associated neurological dysfunction in rats. Administration of AT following TBI has been found to enhance cell proliferation, differentiation and neurogenesis in the dentate gyrus in a brain-derived neurotrophic factor (BDNF) and vascular endothelial growth factor (VEGF)-associated fashion (42, 43). Studies also suggest role of AT in reducing amyloid-β peptide (Aβ 41 and Aβ 43) deposition and thus preventing Aβ-induced microglial activation (44, 45). Atorvastatin treatment has also culminated into quite remarkable outcomes in treating multiple sclerosis (46), experimental autoimmune encephalomyelitis (47), epilepsy (48), depression (49) and spinal cord injury (50, 51). Several evidences underscore the neuroprotective actions exerted by AT treatment as a result of reduction of oxidative stress markers, lipid peroxidation, and restoration of reduced glutathione (52). Additionally, AT-induced abrogation of reactive oxygen species and matrix metalloproteinases (MMPs) upregulation helped in minimizing the inflammation and blood brain barrier disruption (53, 54). In addition to the neuroprotective roles of AT in the setting of neuroinflammatory diseases, evidences for role of AT upon viral infections also deserve special mention. Studies demonstrate anti-viral action of AT against human immunodeficiency virus-1 by interfering with intercellular adhesion molecule-1(ICAM-1) - leukocyte function-associated antigen-1 (LFA-1) interaction (55). Statins have also been reported to play positive role in protecting patients suffering from cardiovascular diseases against influenza-associated signs and symptoms. Lovastatin has also been demonstrated to inhibit dengue (DEN-2 NGC) virus replication in human peripheral blood mononuclear cells (56) and reduced viremia in lovastatin-pretreated animals against DENV-2 infection (57). Lipid lowering actions of atorvastatin, fluvastatin and simvastatin was observed to help restore alteration of cholesterol biosynthesis and transport induced by Hepatitis C virus (HCV) infection and result in improving disease prognosis. This report thus points towards the role of AT as an adjunct therapy in combating HCV infection in patients treated with pegylated-interferon (IFN) and ribavirin (58). Other than some very preliminary studies till date, there are no reports suggesting the possible role of atorvastatin in treating CHPV and JEV infections, and its putative mechanism of action. Therefore, our study provides the first evidence of effect of AT treatment upon CHPV and JEV infections along with a possible mechanism by which propagation of both positive (+ve) and negative (-ve) stranded RNA viruses were abrogated. Our major findings comprise of increased survivability and reduced viral load in brain tissues of AT-treated mouse pups infected with CHPV and JEV. Moreover, *in vitro* as well as *in vivo* experiments also demonstrate that AT treatment lead to down regulation of ER stress-related protein hnRNPC in the context of CHPV and JEV infection. Based on the aforementioned evidences, we hypothesized that AT might be playing role in curtailing CHPV and JEV titer in the neurons by down regulation of cellular hnRNPC. The neuroprotective action of atorvastatin in the face of viral infections was found to be associated with (i) reduced viral reproduction (ii) reduced abundance of proinflammatory cytokines and certain chemokines in the brain (iii) reduced level of cleaved-Caspase 3 and (iv) reduced ER stress-induced RNA binding protein hnRNPC, which was further validated to play a potential role in propagation of viruses.

Earlier report suggests upregulation of proinflammatory cytokines and chemokines by microglial activation in response to CHPV and JEV infection (6, 59). Restricting the development of neuroinflammation is being considered to be a potent strategy in halting the progress of neurodegeneration. We showed that in the atorvastatin-treated groups following CHPV and JEV infections, levels of MCP1 and proinflammatory cytokines plummeted down to values significantly lower than that of CHPV and JEV-infected groups *in vivo* (IL-6, IFNγ in case of CHPV and IL-6, IFNγ, TNF-α) (Fig. 1C and 1D). These data further supported the notion that reduced microglial activation and resulting neuroinflammation is critical in determining the survival rate of the virus-infected pups as demonstrated by their increased survival and delayed appearance of signs in comparison to virus-infected mice not treated with AT (Fig. 1A and 1B). The dosage of AT used (5mg/kg body weight) *in vivo* was safe as evidenced by the effect of the dosage upon body-weights of mouse pups as shown in supplementary figure 1A and 1B. In addition to its role in reducing the severity of inflammatory reaction, atorvastatin was also found to significantly reduce CHPV and JEV propagation in mouse pups. Abundance of viral proteins (G, N and M proteins of CHPV and NS3 of JEV) in AT-treated mice brain in the context of virus infection was found to be significantly lower than that in infected group (Fig. 2A and 2B). Moreover, evidence of action of AT reducing virus-induced neuronal apoptosis has been provided by demonstration of decreased expression of cleaved-caspase-3 fragment in AT treated group with respect to that of only infected pups. This down regulated activation of Caspase-3 corresponds to AT-induced reduction in mortality of infected mice pups (3, 60). Effect of DMSO was also assessed *in vivo* in order to rule out any of its unwanted role upon viral propagation and neuronal apoptosis (Supplementary Fig. 2A and 2B). We also evaluated the effect of virus infections in absence or presence of AT treatment upon ER stress response since earlier reports suggest that stimulus-induced ER stress response can result in various diseases including neurodegeneration (61). Studies conducted by many independent groups demonstrate that infections by flaviviruses like dengue virus, hepatitis C virus and West Nile virus can lead to induction of unfolded protein response (UPR) (62, 63, 64). Work by our group also provides testimony to the fact that JEV exerts detrimental effects upon human neural stem cells and sub ventricular zone (SVZ) of mouse pups via activation of ER-stress characterized by upregulation of endoplasmic reticulum-resident chaperone (GRP78 or Bip), prohibitin (PHB) and heterogeneous nuclear ribonucleoprotein C (hnRNP C1/C2) (37). In our study, expression levels of the three abovementioned proteins were subjected to analysis upon CHPV and JEV infections in absence and presence of AT. Unlike Bip and PHB, AT treatment was found to decrease virus-induced hnRNPC upregulation *in vivo* (Fig. 2A, 2B, supplementary Fig. 2C and 2D). Subsequently, viral titer of CHPV and JV-infected mice brain samples was found to be reduced by treatment with AT upon viral infections as evidenced from the results of plaque assay (Fig. 2C and 2D). Immunohistochemical analysis of CHPV and JEV-infected brain sections revealed upregulated expression of hnRNPC in virus-infected cells. Interestingly, AT treatment was shown to restore the enhanced cellular abundance of both hnRNPC and viral proteins (Fig. 3A and 3B). Since both CHPV and JEV are known to infect neuronal cells, to assess the action of AT upon host hnRNPC and viral propagation in neurons we cultured cortical neuronal cells isolated from mice cerebral cortex followed by immunohistochemical analysis. In line with *in vivo* findings, both CHPV or JEV proteins and hnRNPC expression was observed to be suppresses upon treatment of mouse primary cortical cells with AT (Fig. 5A and 5B).

Taken together, analysis of our *in vivo* experiments’ data reveals that AT treatment in the context of CHPV and JEV infection results in decreased mortality which was also found to be associated with reduced ER stress, CHPV and JEV viral propagation. Our observations from *in vivo* experiments were also further strengthened by similar outcomes in *in vitro* experiments. When AT was analyzed for cytotoxicity, dosages up to 2 µM were found not to exert any significant effect upon viability of neuro2A cells (Supplementary Fig. 3A). Since AT was reconstituted in DMSO, effect of DMSO upon viral propagation and cell viability was also subjected to investigation. No significant changes in virus reproduction and cell viability were observed *in vitro* upon administration of DMSO (Supplementary Fig. 3B and 3C).

Elevated intracellular ROS level in the CHPV and JEV infected Neuro2A cells and inhibition of ROS with the atorvastatin treatment (2 µM) further strengthen our notion related to the infection and ER stress-mediated response (Supplementary Fig. 4A and 4B). We further checked the effect of different dosages of AT treatment upon CHPV and JEV propagation, ER stress-induced hnRNPC abundance and cell viability *in vitro*. AT was found to reduce both CHPV and JEV propagation (as exemplified by G, N and M proteins in case of CHPV and NS3 in case of JEV), hnRNPC and cleaved-caspase-3 expression in a dose-dependent fashion (Fig. 4A and 4B). Moreover, plaque assay analysis demonstrated reduced viral titer from cell supernatant upon treatment with 2 µM atorvastatin when compared to that of virus-infected cells (Fig. 4C and 4D). We also pretreated the Neuro2A cells with thapsigargin and checked the role of atorvastatin treatment (2 µM) directly on the ER stress to get the same response about hnRNPC expression and related apoptosis (cleaved Caspase 3) without virus infection (Fig. 4E).

In the present study, we find that the change in hnRNPC abundance due to CHPV and JEV infections gets restored successfully upon treatment with AT. This has led us to hypothesize that AT might act via host hnRNPC modulation thus culminating into abrogation of viral propagation. According to multiple earlier reports, hnRNPC has been proposed to act as an RNA-binding protein thus enabling the same to interact with 5′ end of poliovirus negative-strand RNA as well as with poliovirus nonstructural proteins. These hnRNPC-viral genome interactions have been reported to promote efficient positive-strand RNA synthesis (38). There are reports suggesting that hnRNPC could accumulate in the cell cytoplasm to support the virus positive strand replication (65–67). As our earlier work suggests direct interaction of hnRNPC with both JEV and CHPV RNA genomes (37), we were interested to assess role of atorvastatin upon recruitment of viral genome into hnRNPC-containing replication assembly *in vivo* and *in vitro*. We conducted RNA Co-IP experiment to find out that AT treatment resulted in significant reduction of CHPV and JEV genome interaction with hnRNPC (Fig. 6A and 6B; Fig. 7A and 7B). Since AT treatment has already been demonstrated to suppress infection-induced hnRNPC upregulation, this reduced binding of viral genome *in vivo* and *in vitro* suggests that AT-induced hnRNPC down regulation thus leads to inefficient viral replication complex formation, hence resulting in reduced CHPV and JEV propagation. Earlier studies suggest that Polio virus genome replication is modulated *in vitro* with overexpression of recombinant hnRNPC or depletion of hnRNPC proteins (38). Similar effects of hnRNPC knockdown upon Dengue virus genome replication have been demonstrated recently thus underscoring the importance of hnRNPC as a positive regulator of RNA-virus genome duplication (68). To further validate our findings, we studied the effect of hnRNPC knockdown upon infection in the presence or absence of atorvastatin (2 µM). Analysis of viral propagation, exemplified by G, N and M proteins for CHPV and NS3 protein representing JEV, exhibited further reduction upon hnRNPC knock down in the presence of AT. These results reinstate the fact that AT-induced hnRNPC downregulation plays a major role in retarding both CHPV and JEV propagation (Fig. 8A and 8B). Consistent with the abovementioned findings, CHPV and JEV viral titer as measured by plaque assay was also observed to be reduced in the presence of AT treatment (Fig. 8C and 8D). Interestingly, overexpression of hnRNPC in neuro2a cells resulted in increased viral RNA, protein synthesis, which was found unaffected by the presence of AT (2 µM). Measurement of viral titer by plaque assay analysis exhibited increased CHPV and JEV propagation as a result of hnRNPC overexpression, irrespective of atorvastatin treatment.

In addition to being a crucial determining factor of positive-strand RNA virus replication (61–64), hnRNPC is also reported to play many important roles in multiple cellular processes via modulation of numerous cellular targets (39, 40). hnRNPC has been documented to increase miR-21 expression thus resulting in downregulation of its target PDCD4 in case of glioblastoma (39). Another report by Eto *et al* suggests that loss of PDCD4 expression helps augment induction of apoptotic response by promoting translation of procaspase-3 mRNA (41). Since both CHPV and JEV are reported to successfully infect neurons thus resulting in their apoptotic death, and AT has been demonstrate to act via hnRNPC-dependent fashion in reducing cell death, we investigated whether AT action results in diminution of neuronal apoptosis in a miR-21 and PDCD4-associated fashion. Both *in vivo* and *in vitro* experiments using Balb C mice and neuro2A cells respectively display a reduction in CHPV/JEV-induced miR-21 upregulation upon atorvastatin treatment (Fig 10A and 10B), thus confirming role of AT in altering miR-21 abundance. Human autopsy brain samples infected with JEV also exhibit increased miR-21 abundance (Fig: 10C). Targeting cellular hnRNPC using hnRNPC-specific esiRNA in CHPV/JEV-infected cells receiving atorvastatin treatment further reduced miR-21 abundance when compared to that in untransfected-cells infected with CHPV/JEV as well as receiving AT treatment (Fig: 10D and 10E) thus pointing towards the causal association in between infection-mediated hnRNPC upregulation and increased miR-21 expression. To further validate the operation of AT treatment-hnRNPC-miR-21 axis in the setting of CHPV and JEV infections, cells infected by CHPV/JEV and receiving AT treatment were transfected with plasmids coding hnRNPC. Plasmid-mediated hnRNPC overexpression in CHPV/JEV-infected cells which also received AT treatment culminated into restoration of miR-21 abundance to a level comparable to that of cells infected with CHPV/JEV (Fig: 10F and 10G). Since atorvastatin treatment leads to reduction in infection-mediated rise in hnRNPC abundance, a significant decline in miR-21 expression was observed. Whereas, restoration of cellular hnRNPC as a result of overexpression helped restore miR-21 upregulation thus confirming role of hnRNPC in regulating miR-21 expression in a positive fashion.

To decipher miR-21’s mechanism of action upon CHPV/JEV infection and its modulation upon atorvastatin treatment, abundance of PDCD4, a target of miR-21 as reported earlier (39), were studied both *in vivo* and *in vitro* during the course of CHPV and JEV infections (Fig: 11A and 11B). PDCD4 expression was shown to be downregulated at protein level in case of both CHPV and JEV infection studies *in vivo* and *in vitro* (Fig: 11A and 11B). Inhibition of miR-21 activity in CHPV and JEV-infected cells resulted in enhanced PDCD4 expression and reduced caspase-3 activation with respect to CHPV/JEV-infected or control anti-miR-transfected cells also infected by CHPV/JEV (Fig: 11C and 11D). Owing to that fact that miR-21 targets PDCD4 expression (39), CHPV and JEV infection-mediated miR-21 upregulation culminated into abolished PDCD4 expression thus eliciting activation of Caspase-3. On the other hand, inhibition of miR-21 by inhibitor resulted in restoration of PDCD4 expression thus abrogating activation of Caspase-3. Taken together, these abovementioned findings prove role of miR-21 in triggering apoptotic response upon infection by CHPV and JEV. In order to find out whether atorvastatin does act as a neuroprotective agent in the context of CHPV and JEV infection by interfering with miR-21 activity, effect of atorvastatin upon miR-21 and PDCD4 abundance was studied following CHPV and JEV infection *in vitro*. CHPV and JEV-induced reduction in PDCD4 expression and enhanced miR-21 abundance was restored upon treatment of infected-neuro2A cells with atorvastatin both *in vitro* and *in vivo* (Fig: 11E and 11F). Since PDCD4 expression acts to antagonize Caspase-3-dependent cell death, atorvastatin-mediated reduced miR-21 expression and enhanced PDCD4 abundance thus might contribute to its neuroprotective role in the context of CHPV and JEV infections, as exemplified by reduced neuronal cell death.

To summarize, our current study provides with the first possible mechanistic details explaining mode of atorvastatin action in combating CHPV and JEV infections. The study suggests that infection-induced ER stress helps in upregulation of hnRNPC which regulates CHPV and JEV propagation in a positive fashion. Moreover, substantial proof has been provided by our work demonstrating role of atorvastatin treatment in abrogating hnRNPC abundance, thus ameliorating viral propagation. In addition to that, atorvastatin-mediated hnRNPC downregulation has also been shown to alter virus-induced miR-21 upregulation. Via suppressing cellular expression of miR-21, atorvastatin results in increased PDCD4 abundance and hence reduced Caspase-3 activation, manifested as diminished neuronal death in response to viral infections. Since atorvastatin has been used as a well-tolerated hypolipidemic drug aimed at combating lipid-related disorders, further clinical assessment of atorvastatin’s efficacy in fighting neurotropic viral infections may provide us with future possible therapeutic strategy in JEV and CHPV endemic regions.

## Acknowledgements

Authors sincerely acknowledge Prof. Anita. Mahadevan, Department of Neuropathology, National Institute for Mental Health and Neurosciences, Bangalore, for providing autopsied human tissue. We are thankful to I. Akbar, K. L. Kumawat and M. Dogra for their technical assistance.

## Supplementary materials

**Supplementary Fig. 1:**
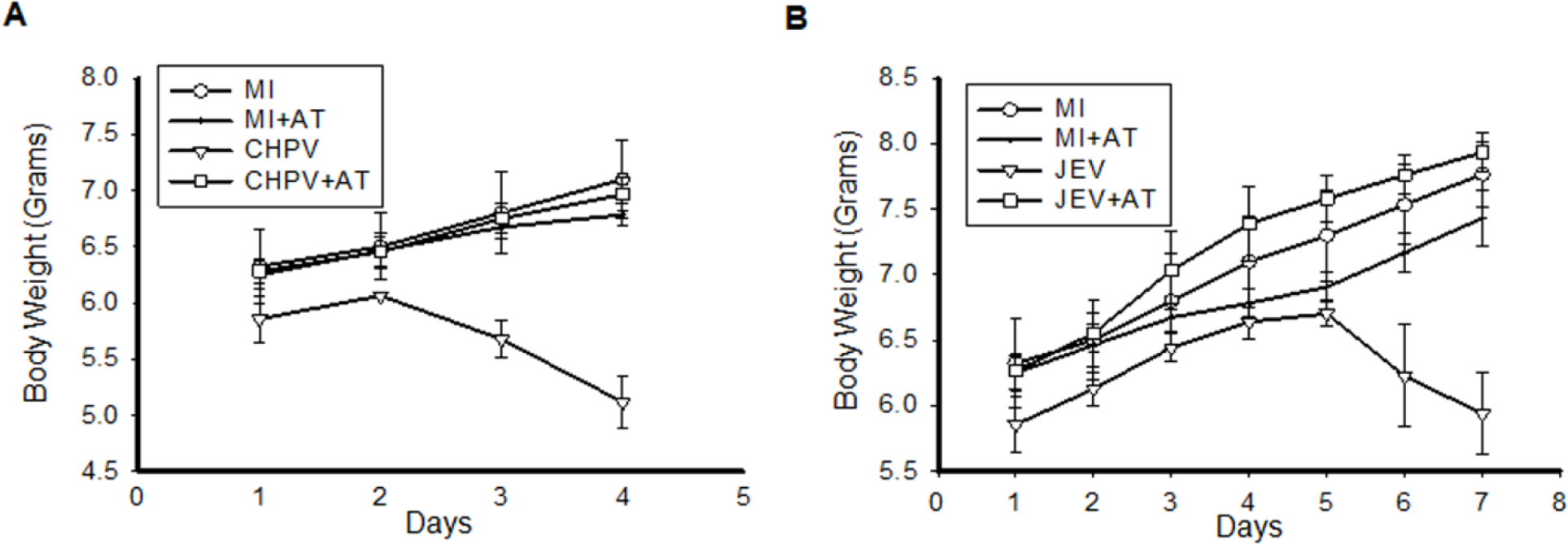
Atorvastatin helped maintain normal body-weight in the context of CHPV and JEV infection. (A) CHPV-infected mice treated once daily with atorvastatin (AT) at a dosage of 5mg/kg-body weight till day 4 following infection, and monitored carefully for any changes in body-weight in comparison to other experimental groups. (n=10 in each group) (*p ≤ 0.001). (B) JEV-infected mice were treated with AT once daily at a dosage of 5mg/kg-body weight till day 7 following infection, and observed carefully for any changes in body-weight with respect to other experimental groups. (n=10 in each group) (*p ≤ 0.001).

**Supplementary Fig. 2:**
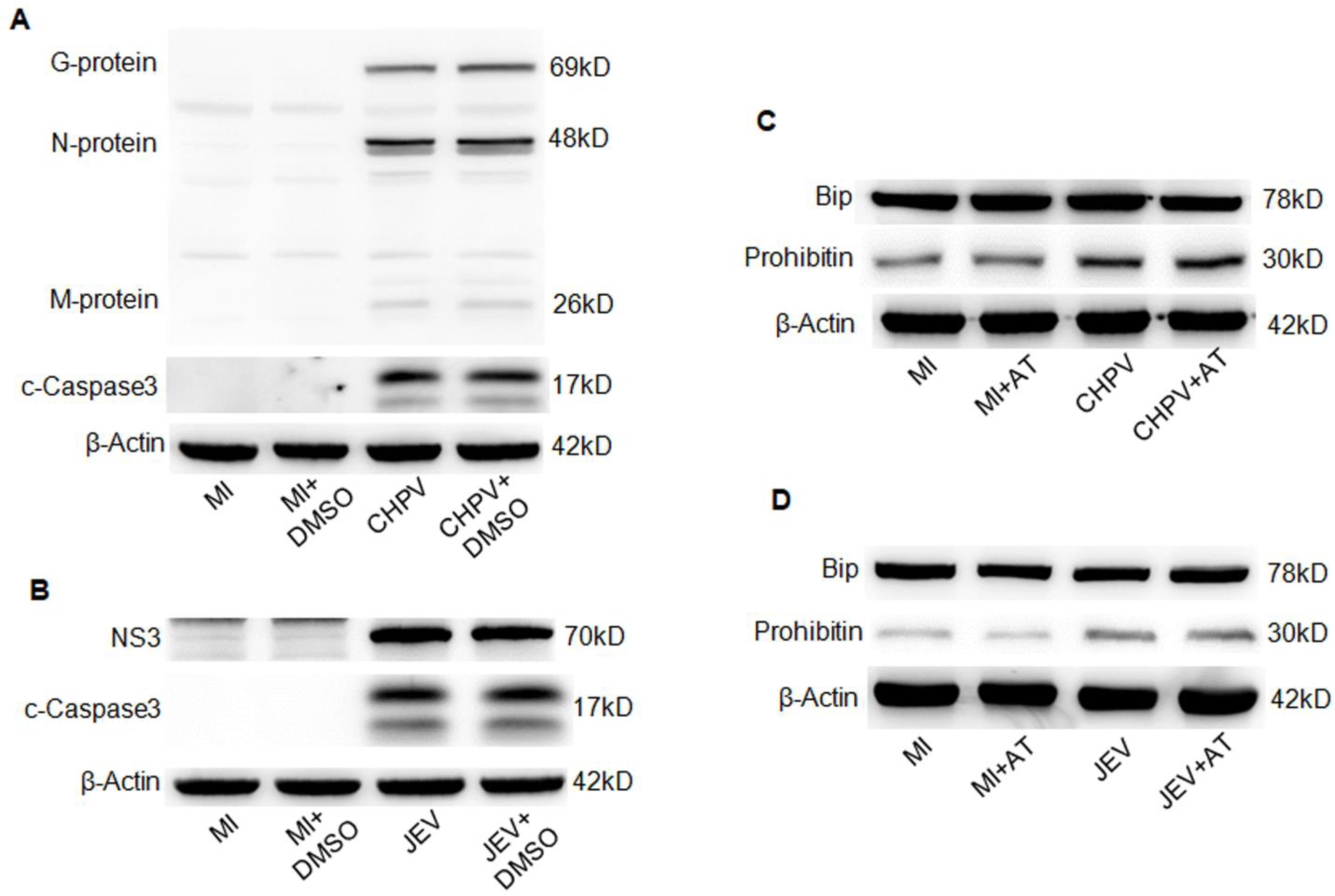
Effect of DMSO upon cellular viability and atorvastatin (AT) upon virus-induced ER stress response. (A) In order to assess the effect of DMSO on viral replication efficiency, four mice groups have been studied here. Mock infected (MI), MI with DMSO, CHPV-infected group and Virus-infected group where CHPV was resuspended in DMSO. Once signs of infection became evident following 3 days of infection, brain samples were isolated followed by homogenization. The samples were then processed for immunoblotting for assessing abundance of three proteins coded by the viral genes, viz. Glycoprotein (G), Nucleocapsid protein (N) and Matrix protein (M) along with cleaved fragment of Caspase-3. β-actin protein expression was used as loading control. (B) (C) Immunoblotting analysis of Bip and prohibitin expression across the four groups of mice (MI, MI with AT, CHPV and CHPV with AT) was performed to study the effect of AT treatment upon ER stress. Followed by 3 hours of infection with CHPV, infected-mice were treated with AT. AT treatment was provided to mock-infected mice and was subjected to analysis of ER stress. β-actin was used as loading control for all experimental groups. (D) Similar to figure shown in (C), differences in expression of Bip and Prohibitin of JE virus-infected sample in the absence and presence of AT was studied using immunoblotting. β-actin was used as loading control for all experimental groups.

**Supplementary Fig. 3:**
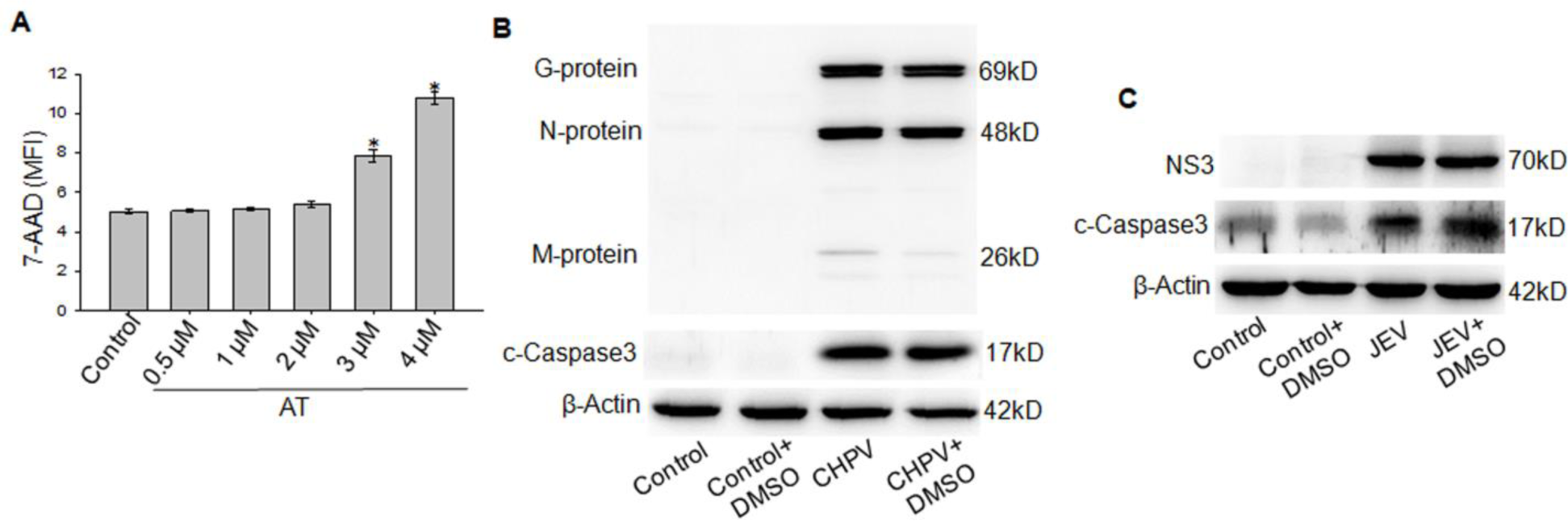
*In vitro* AT Dose-determination and evaluation of effect of DMSO upon infection severity and cell viability. (A) Cell viability of Neuro2A in response to different dosages of atorvastatin (0.5, 1, 2, 3, 4 µM) was measured by staining cells with 7-AAD followed by determination of mean fluorescence intensity (MFI) of 7-AAD in a flow cytometer (*p ≤ 0.001). Data Values are represented as mean ± SD of three independent experiments. (B and C) Effect of DMSO upon CHPV and JEV infection in neuro2A cells were analysed by assessing expression of CHPV (B) or JEV (C)-coded proteins in presence and absence of DMSO. DMSO was added to cell culture followed by 2-hour incubation with virus, and maintained in the maintenance media. Antibodies against G, N, and M proteins of CHPV and NS3 protein of JEV were used for immunoblotting analysis. Cell viability in response to DMSO was also studied by examining the abundance of cleaved-fragment of Caspase-3.

**Supplementary Fig. 4:**
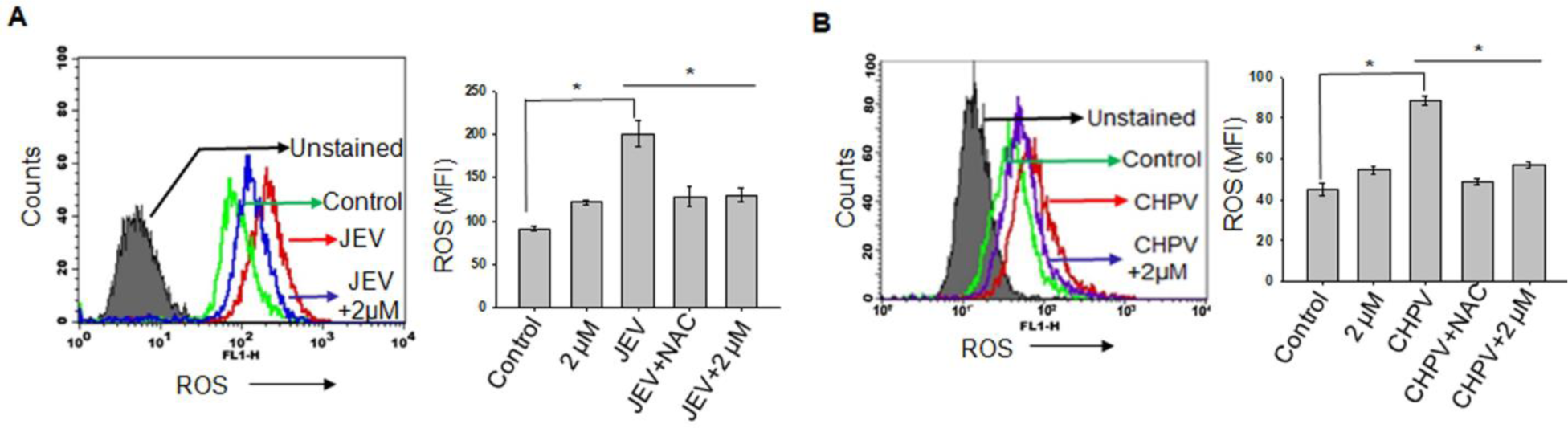
AT treatment resulted in abrogation of CHPV/JEV-induced generation of cellular reactive oxygen *in vitro*. (A) Neuro2A cells were incubated with JEV for 2 hours followed by 3 hours of atorvastatin treatment (2 µM). Collected cells were then subjected to DCFDA staining and analysed by flow cytometry. (B) Neuro2A cells were incubated with CHPV for 2 hours followed by AT treatment as stated in (A). Treated cells were then stained with DCFDA and ROS generation was assessed using flow cytometry. Cells were treated with N-acetyl cysteine (NAC 10 µM)) for 2 hours prior to JEV/CHPV infections and used as negative control. Histogram showed represents one of the three independent experiments and the graph represented is mean ± SD of MFI of three independent experiments (*p ≤ 0.001).

